# A theory of actions and habits: The interaction of rate correlation and contiguity systems in free-operant behavior

**DOI:** 10.1101/807800

**Authors:** Omar D. Perez, Anthony Dickinson

**Author notes:** Correspondence concerning this article should be addressed to Omar D. Perez, 1200 East California Boulevard, Pasadena, California, CA91107.(c) 2020, American Psychological Association. This paper is not the copy of record and may not exactly replicate the final, authoritative version of the article. Please do not copy or cite without authors’ permission. The final article will be available, upon publication, via its DOI: 10.1037/rev0000201.

## Abstract

Contemporary theories of instrumental performance assume that responding can be controlled by two behavioral systems, one goal-directed that encodes the outcome of an action, and one habitual that reinforces the response strength of the same action. Here we present a model of free-operant behavior in which goal-directed control is determined by the correlation between the rates of the action and the outcome whereas the total prediction error generated by contiguous reinforcement by the outcome controls habitual response strength. The outputs of these two systems summate to generate a total response strength. This cooperative model addresses the difference in the behavioral impact of ratio and interval schedules, the transition from goal-directed to habitual control with extended training, the persistence of goal-directed control under choice procedures and following extinction, among other phenomena. In these respects, this dual-system model is unique in its account of free-operant behavior.

## Introduction

Instrumental action instantiates a unique reciprocal relationship between the mind and the world. Through instrumental learning we bring our representations of the consequences or outcomes of our actions into correspondence with the causal relationships in the world, whereas through instrumental action we bring the world into correspondence with the representations of our desires. However, this reciprocity assumes that instrumental behavior is goal-directed in the sense that it is based upon an interaction between a belief about the causal response-outcome relationship and a desire for the outcome (Dickinson and Balleine, 1994; Heyes and Dickinson, 1990). Over the last forty years a wealth of evidence has accumulated that not only are humans capable of goal-directed action in this sense, but so are other animals.

The canonical assay for the goal-directed status of instrumental behavior is the outcome revaluation procedure, which we shall illustrate with an early study by Adams and Dickinson (1981). They initially trained hungry rats to press a lever to receive either sugar or grain pellets with the alternative reward or outcome being delivered freely or non-contingently. The lever was then withdrawn and a flavor aversion was conditioned to one type of pellet by pairing its consumption with the induction of gastric malaise until the rat would no longer eat this type of pellet when freely presented. The purpose of this outcome devaluation was to remove the rat’s desire for this type of pellet, while maintaining the desirability of the other type. If lever-pressing was mediated by knowledge of the causal relationship with the pellet outcome, devaluing this outcome should have reduced the rat’s propensity to press when the lever was once again presented relative to the level of responding observed when the non-contingent pellet was devalued. This is exactly the result they observed (Adams and Dickinson, 1981). More recently, the finding has also been documented in both humans (Valentin *et al.*, 2007) and monkeys (Rhodes and Murray, 2013).

It is important to note that this test is conducted under extinction with the delivery of the outcome suspended; any devaluation effect should therefore reflect knowledge acquired during training rather than during the test itself. A decrease in responding without further experience with the outcome produced by the instrumental response is evidence that the animal represents both the response-outcome contingency and the current outcome value so that the integration of these two representations is reflected in test responding, and it is for this reason that we characterize the responding as goal-directed (Dickinson and Perez, 2018).

Although research on the brain systems supporting goal-directed behavior has advanced during the last 20 years (for a review, see Balleine, 2019b; Balleine and O’Doherty, 2010), the nature of the psychological processes underlying the acquisition of response-outcome knowledge remains relatively under-studied. This is in part because the psychology of learning has focused on the Pavlovian paradigm for the last 50 years or so given the greater experimental control afforded by such procedures. This research has generated a rich corpus of associative learning theories, all of which assume that learning is driven, in one way or another, by prediction errors (for a review, see Vogel *et al.*, 2004). In the case of Pavlovian learning, these errors reflect the extent to which the conditioned stimulus fails to predict to the occurrence (or non-occurrence) of the outcome. In the most straightforward of these theories, the larger the prediction error on a learning episode the less predicted is the outcome and the greater is the change in associative strength of the stimulus. As a consequence, the prediction error is reduced appropriately on subsequent, congruent learning episodes (Rescorla and Wagner, 1972). Based on the idea that Pavlovian associative learning is controlled by prediction errors and the multiple phenomena that parallel those found in instrumental learning, Mackintosh and Dickinson (1979) suggested such errors play an analogous role in both types of learning processes.

Over the last decade or so, goal-directed learning has become increasingly couched in terms of computational reinforcement learning (RL). According to this approach (Daw *et al.*, 2005; Maia, 2009; Sutton and Barto, 1998), goal-directed behavior is controlled by model-based computations in which the agent learns a model of the state transitions produced by the instrumental contingencies and the value of each of the experienced states. At the time of performance, the agent searches the model to estimate the value of each of the actions available, and chooses the one that maximizes the outcome probability over a number of episodes of acting on the environment. Critically, what determines the value of each action in each state (or, alternatively, the probability of choosing each of the available actions in each state) is the probability that a rewarding outcome will be received given that the action is performed. Similarly, theories based on Bayesian inference (Solway and Botvinick, 2012) also deploy the conditional probability of an outcome given an action as a critical variable.

Whatever the differences between the associative, RL, and Bayesian accounts of goal-directed action, these approaches share the assumption that the probability of a rewarding outcome is a primary determinant of instrumental goal-directed action. The reward probability directly determines the strength of the response-outcome association according to associative theory (Mackintosh and Dickinson, 1979) and the estimated value of an action in the case of RL and Bayesian theories. For all of these approaches, instrumental performance should be directly related to the probability of an action leading to a rewarding outcome. However, ever since the initial studies of instrumental outcome revaluation using free-operant schedules we have known that reward probability is unlikely to be the primary determinant of goal-directed control.

### Ratio and interval contingencies

In contrast to the successful demonstration of devaluation reported by Adams and Dickinson (1981), prior investigations of goal-directed free-operant behavior had all trained rats to press the lever on a variable interval (VI) contingency between the response and the outcome and were uniformly unsuccessful (Adams, 1980; Holman, 1975; Morrison and Collyer, 1974). This class of schedule models a resource, such as nectar, that depletes when taken and regenerates with time. In practice, a VI schedule specifies the average time interval that has to elapse before the next outcome becomes available. In contrast, Adams and Dickinson (1981) used a variable ratio (VR) schedule, which models foraging in a non-depleting source so that each action has a fixed probability of yielding an outcome independently of the time elapsed since the last outcome was obtained.

In an experimental analysis of the ratio-interval contrast, Dickinson et al. (1983) used a yoking procedure to match the outcome probability on the two schedules. In one pair of groups, the master rats were trained in an interval schedule, whereas the yoked animals were trained on ratio schedules with outcome probabilities that matched those generated by the master rats. Figure 1 shows their results. As can be appreciated, in spite of the fact that the outcome probability per response was matched between the groups, outcome devaluation reduced performance of the ratio-but not the interval-trained group, suggesting that ratio training more readily establishes goal-directed control than interval training. This conclusion was reinforced when the outcome rate was matched by yoking the rates of the interval-trained rats to those generated by master ratio-trained animals. Again, ratio-, but not interval-trained animals, were sensitive to outcome devaluation. As the interval-trained rats pressed at a lower rate than the ratio-trained animals, goal-directed control was observed in the ratio-trained group even under a lower outcome probability experienced by those rats. The higher levels of performance produced by ratio training under matched reward probabilities (Catania *et al.*, 1977; Pérez *et al.*, 2016; Pérez and Soto, 2020) and the impact of the training schedule on the outcome devaluation effect (see Gremel and Costa, 2013; Hilario *et al.*, 2012; Wiltgen *et al.*, 2012) have now received extensive replication.

**Figure 1.**
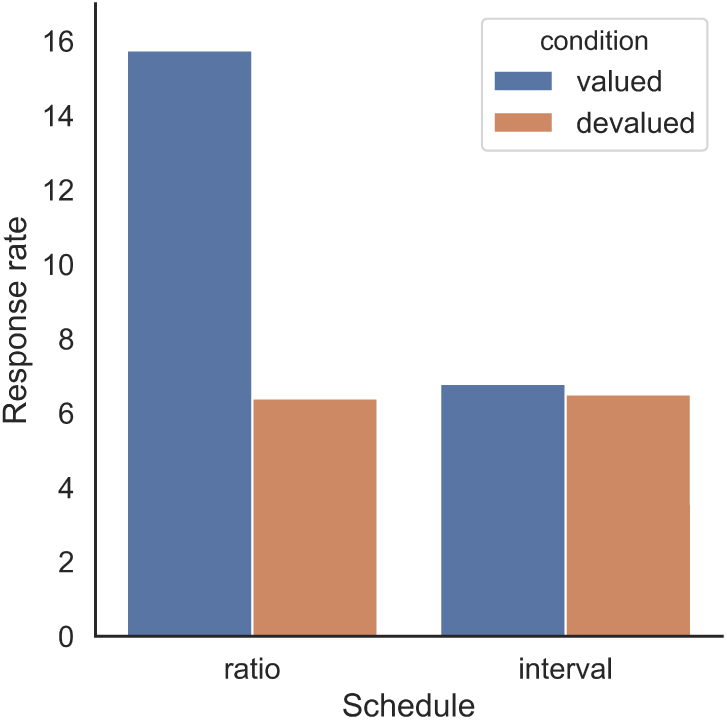
Results obtained by Dickinson and colleagues (1983) in an experimental analysis of behavioral performance and control (i.e., sensitivity to outcome devaluation) in rats. The ratio group was yoked to the interval group by matching the reward probability experienced by the interval-trained rats. Goal-directed control of behavior was only evident in the ratio-trained animals.

The claim that ratio training more readily establishes goal-directed control than does interval training when outcome probabilities (or rates) are matched finds further support by a study of the acquisition of beliefs about the effectiveness of an action in causing an outcome. Reed (2001) trained human participants on a fictional investment task in which pressing the space-bar on the keyboard acted as the instrumental response. Ratio training uniformly yielded higher judgments of the causal effectiveness of the key-press in producing the outcome than did interval training both when the probability or rate of the outcome was matched by within-participant matching.

### Two properties of reward schedules

Our review raises the issue of the critical feature of the two types of schedule that determines the relative sensitivity of ratio and interval performance to outcome revaluation. There are two properties that distinguish the contingencies. The first is that variable interval contingencies differentially reinforce pausing between responses or, in the operant conditioning jargon, long inter-response times (IRTs). Indeed, having performed a response and collected the outcome if available, the longer that the agent waits before performing the next response, the more likely it is that the resources will have regenerated so that the next response performed will be rewarded with the outcome. Figure 2a illustrates the relationship between the seconds elapsed since the last response and the probability of the next response being rewarded for different parameters of a random interval (RI) schedule under which there is fixed probability of an outcome becoming available in each second. As can be appreciated, the probability of reinforcement increases monotonically with the time between responses at a faster rate with shorter programmed intervals between outcomes. In contrast, this probability is independent of the pause to the next response under a ratio contingency that arranges a fixed probability of reward that is independent of the time elapsed since the last response^1^.

**Figure 2.**
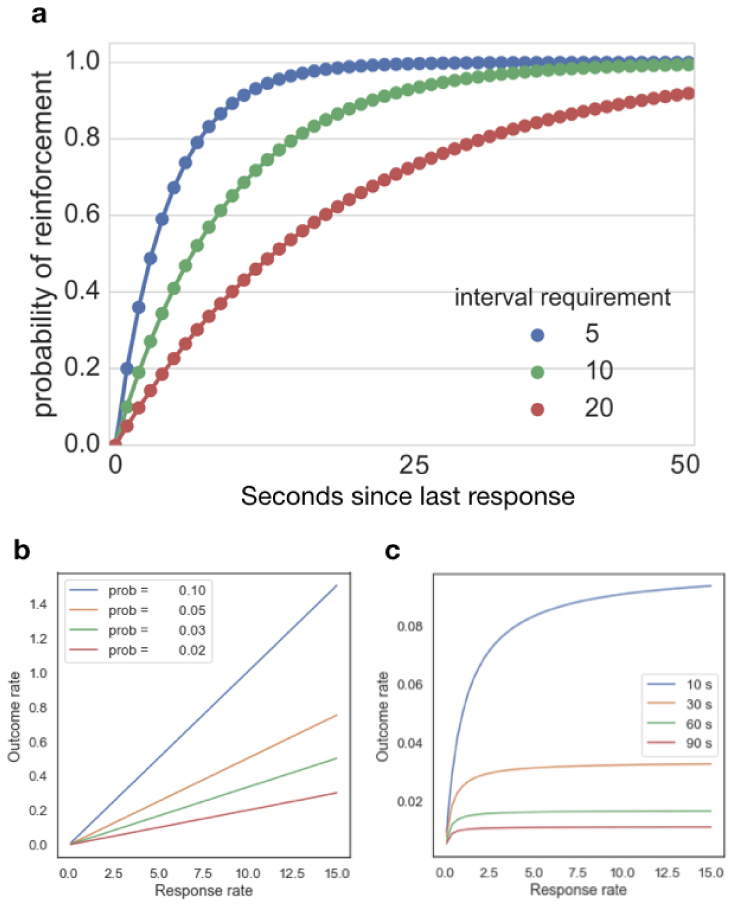
Different properties of reward schedules. (a) Probability of obtaining and outcome after a pause between responses for different programmed inter-reinforcement intervals under a random interval (RI) schedule. (b) Functional relationship between response rates (in responses per minute) and outcome rates (in rewards per minute) for ratio schedules with different outcome probabilities. (c) Functional relationship between response rate (in responses per minute) and outcome rates (in rewards per minute) for interval schedules under different interval parameters.

It is unlikely that this feature of interval contingencies is the key factor that reduces sensitivity to outcome revaluation because when an animal is trained with a choice between two interval sources yielding different outcomes as opposed to a single interval source, performance is highly sensitive to outcome devaluation (i.e., remains goal-directed). Kosaki and Dickinson, for example, trained their hungry rats with a choice between pressing two levers (group *choice*), one yielding grain pellets and the other a sugar solution, both on RI schedules (Kosaki and Dickinson, 2010). In spite of of giving their rats extended training, devaluing one of the outcomes reduced performance of the corresponding response on test. This goal-directed control contrasted with the insensitivity to outcome devaluation following matched training with a single response. This second, *non-contingent* group of rats was trained with a single lever so that pressing yielded one of the outcomes on the interval schedule with the other being delivered at the same rate but independently, or non-contingently of the instrumental response. In contrast to the goal-directed control observed in the choice group, lever pressing during the test was unaffected by whether the contingent or non-contingent outcome had been devalued. As the target responses were both trained under identical interval schedules, both of which should have differentially reinforced long IRTs, it is not clear why choice versus single response training should affect the degree of goal-directed control if IRT reinforcement is the critical factor affecting sensitivity to outcome revaluation under interval schedules.

The second distinction between ratio and interval contingencies relates to their response-outcome rate feedback functions, which are mathematical descriptions of the empirical relationship between response rates and outcome rates (Baum, 1973; Baum, 1992; Pérez *et al.*, 2016; Reed, 2007; Soto *et al.*, 2006). Figure 2 (panels B and C) presents the feedback functions for typical ratio and interval schedules, respectively. Under a ratio contingency, the outcome rate rises linearly with increasing response rate, with the slope of the function decreasing systematically as the ratio parameter increases. The feedback function for ratio schedules can be described by a linear function of the form *R* = *nB*, where *R* is the outcome rate and *B* the response rate performed by the agent. The parameter *n* represents the inverse of the ratio requirement (1/*ratio*), or, equivalently, the outcome probability per response that the particular ratio schedule programs. By contrast, the feedback function for an interval schedule is nonlinear with the outcome rate rising rapidly with increases in response rates when the baseline response rate is low and reaching an asymptote as soon as the response rate is higher than the rate at which the outcomes become available (Baum, 1992; Prelec, 1982). At this point, variations in response rates do not have an effect in the obtained outcome rate ^2^.

When Baum first proposed a correlational version of the Law of Effect, he (1973) suggested that the difference in performance between ratio and interval schedules when outcome probabilities or rates are matched could be explained by their different feedback functions. To analyze this empirically, he proposed that the feedback function could be captured by the linear correlation between the response and outcome rates, and that it should be higher for ratio contingencies at intermediate to fast rates of responding. This idea led Dickinson and his colleagues (Dickinson, 1985; Dickinson *et al.*, 1995; Dickinson and Perez, 2018; Pérez *et al.*, 2016; Pérez *et al.*, 2018; Pérez and Soto, 2020) to argue that response-outcome learning is driven by the response-outcome rate correlation experienced by the agent: the greater the experienced rate correlation, the stronger is goal-directed control.

In the next section, we develop a computational mnemonic-based system for goal-directed action that yields a response strength on the basis of the experienced response-outcome rate correlation. We then report simulations of the sensitivity of this response strength to some of the major variables of free-operant schedules. In order to accommodate the fact that in many cases free-operant behavior is insensitive to outcome devaluation, or in other words behaviorally autonomous of the current value of the training outcome, we then integrate this goal-directed system with a habit system based on a reinforcement learning algorithm to generate a dual-system model of free-operant behavior. Again, we examine the performance of this model against variations in the amount training, the availability of choice, and under extinction. Finally, we discuss additional predictions of the model and processes that impact on free-operant behavior but that lie outside its scope.

Unlike most models of instrumental action in the computational RL literature, we evaluate our model solely against free-operant behavior. The reasons for this choice are two-fold. As we have already pointed out, the empirical genesis for rate correlation theory lies with the impact of ratio and interval contingencies on behavior, which is essentially a property of free-operant contingencies. Second, and more importantly, free-operant procedures are intrinsically action-rather than stimulus-oriented in that they engage the spontaneous emission of actions in a relatively constant stimulus environment. Moreover, we have good evidence that free-operant actions on a manipulandum, such as a lever, are directly sensitive to the instrumental response-outcome contingency as assessed by the canonical bidirectional conditioning criterion (Dickinson *et al.*, 1996). By contrast, model-based RL is primarily structured for a discrete-trial choice paradigm, such as the two-step choice task (Daw *et al.*, 2011; Doll *et al.*, 2012), in which typically a human agent is instructed to choose between two or more localized visual stimuli. Although the target action is a nominal state variable within a RL model, it is not clear that the agent actually represents the action as an element of the task contingency which, within our theoretical framework, is a criterion for goal-directed status. Alternatively, the agent may simply use the action as a vehicle for registering a stimulus preference while encoding the choice task solely in terms of stimulus-outcome or stimulus-stimulus contingencies.

### Rate Correlation Theory

Baum (1973) illustrated the empirical application of his approach to the Law of Effect by dividing the time-line in an experimental session into a number of successive time samples and displayed the rate correlation by plotting the number of responses in each sample against the number of outcomes in that sample. In the present approach, we develop rate correlation theory in terms of psychological processing by assuming that the agent computes the rate correlation at a given point in time by reference to the contents of a short-term memory. This memory is segmented into a number of time samples spanning the recent past, in each of which is recorded the number of responses and outcomes that occurred in that time sample. When the current time sample expires, the system computes the current rate correlation across the contents of the short-term memory.

Figure 3a illustrates a schematic representation of the time-line divided into different samples in the short-term memory of our model. At the end of each sample, the number of responses and outcomes in that sample is registered in the memory and the content of the short-term memory is recycled. Given that the memory has a limited capacity, for simplicity we assume that this recycling involves not only the registration of the contents of the next sample but also the erasure of the oldest sample in memory. For illustrative purposes, Figure 3a displays a memory of four samples; that is, the initial memory cycle involves the first four samples, the second memory cycle involves the second to fifth samples, and so on. In general, if the memory size of our agent is *N*, cycle *k* involves the deployment of the contents of memory from samples 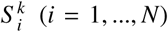. In the simulations shown in this paper, we assume that the memory size is the same for all subjects.

**Figure 3.**
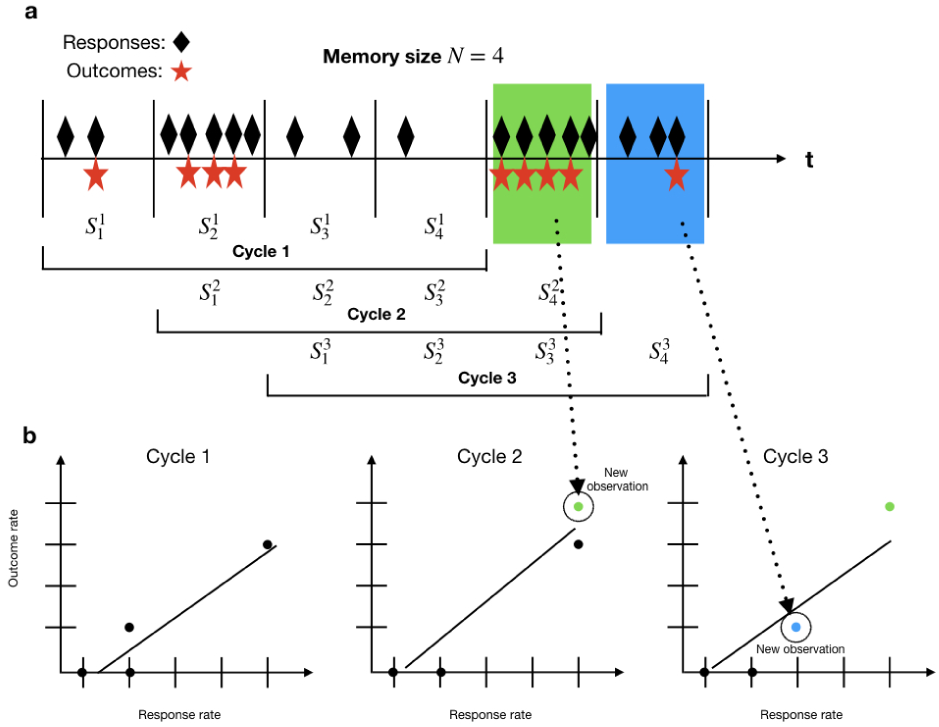
Memory model for a rate-correlation approach to instrumental actions. (a) In this simplified illustration, each memory cycle is composed by four time-samples. The romboids represent response events and the outcomes are represented by red stars. (b) Different experienced rate correlations computed for each of the memory cycles exemplified in (a). The arrows indicate the new observations added at each memory recycle.

Following each mnemonic recycle, the agent estimates the response-outcome rate correlation based upon the current contents of the memory. We assume that the agent computes a standard Pearson correlation coefficient which, in psychological terms, accounts for the agent’s *experienced* linear relationship between the action and outcome rates in the cycle under a given reinforcement schedule. More formally, if *B*_*i*_ and *R*_*i*_ represent, respectively, the number of responses and outcomes in the *i − th* sample within memory cycle *k*, then each sample can be understood as an ordered-pair (*B*_*i*_, *R*_*i*_), *i* = 1, …*N*, from which the agent computes the rate correlation *r*_*k*_ by

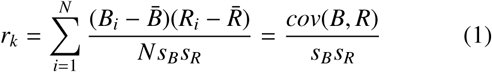

where *cov*(*B, R*) is the covariance between *B* and *R*, 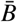 and 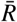 the average responses and outcomes per sample in the cycle, and *s*_*B*_ and *s*_*R*_ the standard deviations of *B* and *R*, respectively.

We propose that goal-directed strength *g* is a direct function of the rate correlation experienced by the agent. To determine the probability of responding *p*, the simplest model would assume that responding during the following cycle *k* + 1 is a function of the rate correlation computed on the basis of the memory contents at the last cycle, that is, *p*_*k*+1_ = *g*_*k*_ = *r*_*k*_. However, there are two concerns about this simple algorithm. First, the algorithm is sensitive solely to the currently experienced rate correlation and so gives no weight to prior experience. This would imply that the strength of the goal-directed system should be immediately affected by abrupt changes in the response-outcome contingency, in contrast with the current empirical evidence. Second, if at a given cycle the memory contains no responses and/or no outcomes, the agent cannot compute a response-outcome rate correlation. From a psychological standpoint, therefore, it is reasonable to assume that the agent relies on both the current and prior experience to determine responding for the next memory cycle. To determine *p*_*k*+1_, therefore, we assume that both the rate correlation experienced in cycle *k* (*r*_*k*_) and the average rate correlation experienced up to cycle 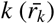 have an effect on the level of responding on the following cycle, with the effect of previous cycles being discounted with time. Under this model, and assuming that there is at least one response and outcome event in memory, the agent can track the average rate correlation 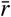 by updating a running estimate according to

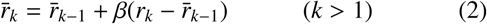

with *β* = 1/*k* as the learning rate in each cycle and 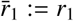, by definition. To determine responding for the following cycle, our agent jointly deploys this average rate correlation with the current experienced rate correlation as follows:

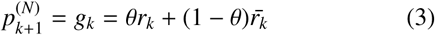

where 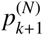 is the probability of responding in the new sample of the following cycle, denoted by 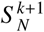, and *θ* is a weighting parameter that represents the importance of the current rate correlation on responding for the next cycle. If *θ* = 1, the agent would put all the weight on the current cycle; if *θ* = 0, its responding would be driven only by the average experienced rate correlation; other values of *θ* will give intermediate degrees of importance to the history of rate correlation on current performance. As noted before, if outcome delivery is suspended under extinction or, more generally, if neither responses nor outcomes are represented in memory, the agent cannot update the estimation of the rate correlation and therefore relies on the last estimation until at least one response and one outcome are again experienced and stored in memory.

It is important to note that the deployment of the average and current rate correlation experienced by the agent determines responding for the next cycle in memory, which only differs with the current cycle in one time sample. The agent responds with strength 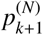 throughout this new sample; the value of *p* is only updated at the end of the cycle according to Equation 3. To simplify the notation, in what follows we ignore the superscript *N* and refer to the response probability of cycle *k* simply as *p*_*k*_, on the understanding that it refers to the response rate in the new sample in memory.

### Simulations of a rate-correlation theory

We first investigated the robustness of the correlation coefficient in this model with respect to variations in the sample duration parameter. To this end, we probed the effect of varying the sample duration between 10 and 120 s on the rate correlation generated by random ratio (RR) 5-to-50 and RI 5- to-90 s schedules with response rates varying between 30 and 150 responses per minute. These two types of schedules assign, respectively, a probability of an outcome being delivered for each response and a probability of the outcome becoming available in each second. In the case of the RI schedule, once the outcome was available, it remained so until collected by a response.

For the range of response rates that we tested, the rate correlation was not significantly affected by the size of the sample chosen (see Supplemental Material), and so we chose a value of 20 s for the time samples in memory, primarily to limit the total duration of the agent’s memory to a few minutes. As the simulations were run with a memory size of 20 samples, the total memory size of each agent was 400 s. For simplicity, we also limited agents’ performance to a maximum of 60 responses per min (i.e. a maximum of 1 response per second) by arranging for the probability of a response in each second to be the value of *g* computed at the recycle that started the current time sample. In what follows, we set *θ* = .5, but note that the same pattern of results hold for the other values of *θ* tested in our simulations.

### Ratio-interval effects

As already noted, our initial reason for investigating the role of rate correlation in goal-directed learning arose from the fact that ratio schedules establish responding that is more sensitive to outcome devaluation than does interval training even when the outcome probability (or rate) is matched by yoking (Dickinson *et al.*, 1983; see Figure 1).

Within our rate correlation theory, *g* is the agent’s learned representation of the strength of the causal relationship between the action and the outcome, which underlies goal-directed behavior. However, as *g* also determines the probability of responding, our theory predicts concordance between judgments of the causal strength of the response-outcome relationship and the rate of responding emitted by the agent.

The most direct evidence for such concordance comes from the study by Reed (2001), who reported the instrumental performance of human participants on ratio and interval schedules with matched outcome probabilities. Not only did he find that ratio training yielded higher causal judgments of the effectiveness of the action in producing the outcome, but also that participants performed at a higher rate under ratio than under interval training. To investigate whether a rate-correlation model could reproduce these data, we simulated training on a master RI 20-s schedule (the temporal parameter employed by Reed (2001) in his human study) and then used the outcome probability generated by each master subject to determine the parameter for a yoked subject trained under a ratio schedule. The initial response rate during the first cycle was 10 per min, and we trained the simulations across 3 sessions, each of which terminated after 13 outcomes, in an attempt to match the training received by the participants in Reed’s (2001) experiment. Figure 4 shows the data obtained by Reed (left panel) and the simulations produced by the rate-correlation model of the response strength, *g*, during the last 50 cycles and averaged across 100 replications of each simulation. As can be appreciated in the right panel of Figure 4, the model generated lower goal-directed response strength values following interval than ratio training, in agreement with Reed’s results.

**Figure 4.**
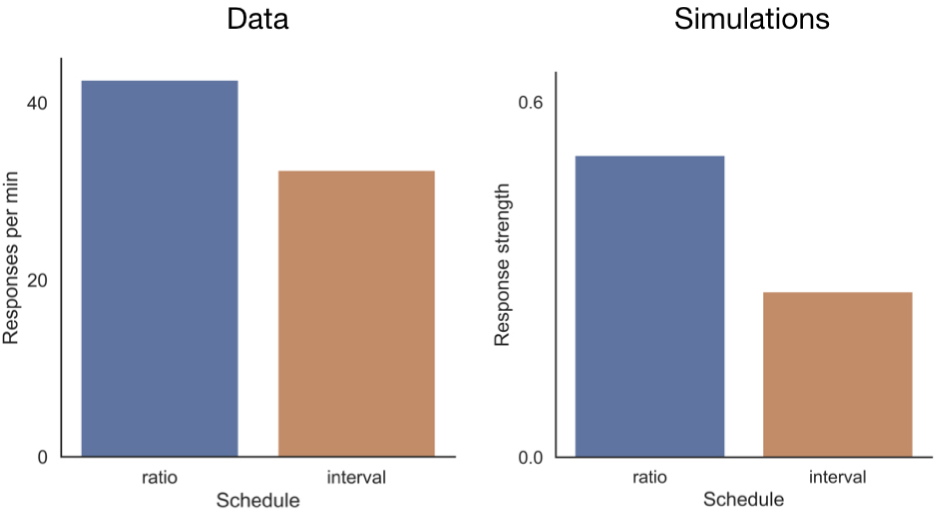
Simulations of a rate correlation model for ratio and interval schedules with matched outcome probabilities. The left panel shows the data obtained by Reed (2001) in a human causal judgment experiment. The right hand panel show the simulations produced by a rate correlation model.

### Outcome probability

Having established that rate correlation theory can reproduce the ratio-interval difference, we investigated whether our model could capture the general effects of the major variables determining free-operant performance. From these simulations we report the response strength, *g*, during the last 50 cycles from the 2000 cycles of each simulation averaged across 100 replications of each simulation. The response rate for the first memory cycle was again set to 10 per min.

We have already noted that both associative and model-based RL theories of goal-directed behavior assume that instrumental learning is driven by reward-prediction error, which in turn implies that instrumental performance should be determined by outcome probability (Mackintosh and Dickinson, 1979; Sutton and Barto, 1998). This prediction was confirmed empirically by Mazur in a free-operant procedure (1983). He trained hungry rats to press a lever on an RR schedule under different ratio requirements (or the inverse of reward or outcome probability). To ensure that the motivational state of the animals was kept relatively constant, Mazur scheduled a limited number of food outcomes per session in an open economy ^3^. In addition, to assess performance only during periods of engagement in the instrumental action, Mazur removed the outcome handling time by assessing the rate following the first lever press after an outcome delivery. The left panel of Figure 5 shows a relevant selection of the response rates obtained by Mazur.

**Figure 5.**
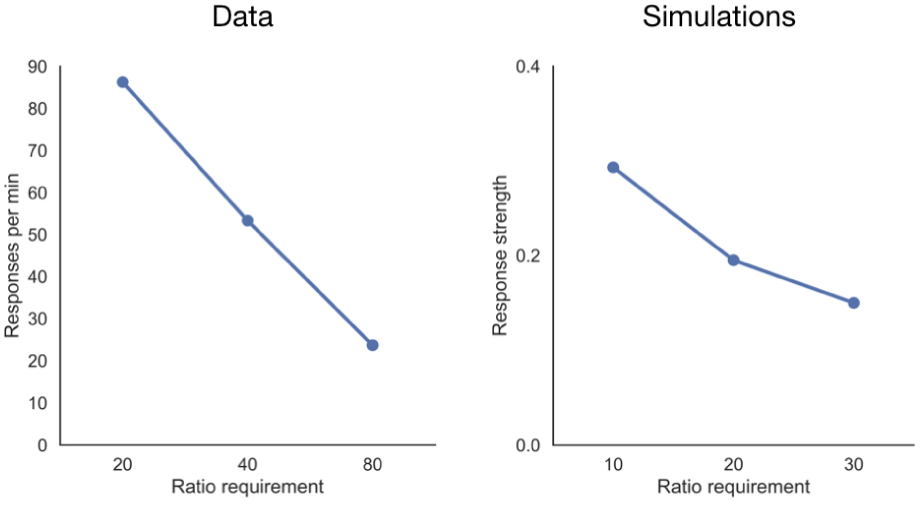
Simulations of rate correlation models for ratio training with different outcome probabilities (the inverse of the ratio requirement). The left panel shows the results obtained by Mazur (1983) in a within-subject study in rats. The right panel shows the simulations of a rate correlation model for a similar design.

To investigate the response rates generated by a rate correlation model when the outcome probability was varied, we replicated a similar design by simulating performance on RR schedules with ratio requirements varying between 10 and 30. Figure 5 shows that the likelihood of responding decreased systematically when the outcome probability was reduced by increasing the ratio parameter, correctly predicting the pattern of results obtained by Mazur in his parametric investigation of ratio performance in rats.

### Outcome rate

As discussed previously, Herrnstein and his colleagues have argued that instrumental performance on interval schedules is systematically related to the outcome rate programmed by the schedule, such that longer programmed intervals between reinforcers should bring about lower performance than shorter ones (Herrnstein, 1969). This prediction has been confirmed multiple times and in different species. One example, shown in the left panel of Figure 6, was provided by Bradshaw et al. (1981), who trained hungry rats to lever press for milk and found a systematic decrease in the response rates as the interval parameter was increased. A selection of their results for intermediate intervals are shown in the left panel of Figure 6. To match the conditions of this experiment, we simulated training under RI schedules with different outcome rates by varying interval parameters between 30 and 90 s. As the right panel of Figure 6 shows, all values produced a systematic decrease in responding as the outcome rate was reduced by increasing the temporal parameter of the interval schedule, replicating the pattern of results obtained by these authors.

**Figure 6.**
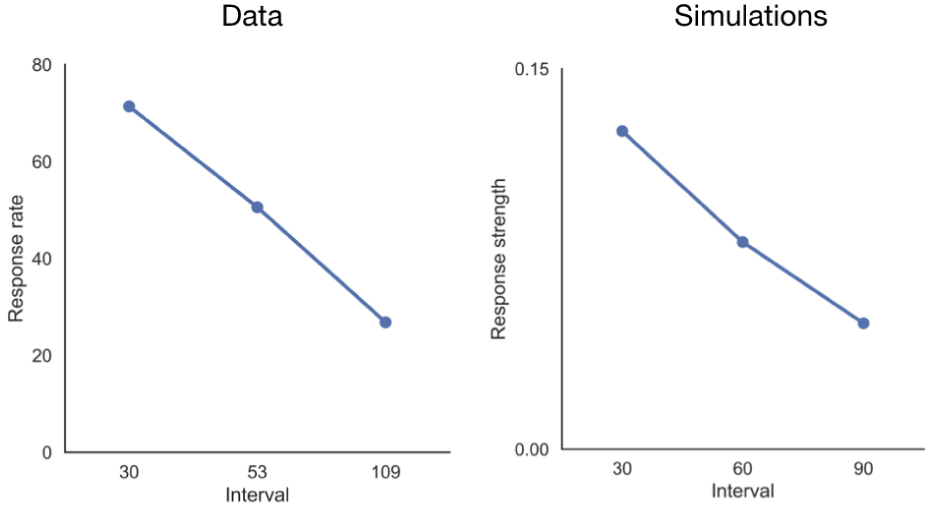
Simulations of a rate correlation model for interval schedules with different interval parameters. The left panel shows the results obtained by Bradshaw et al. (1981) in rats. The right panel shows simulations of a rate correlation model using parameters similar to the ones used in their study.

### Outcome delay

In addition to discussing ratio and interval performance, Baum (1973) noted that his correlational Law of Effect anticipated a deleterious impact on the acquisition and maintenance of instrumental responding when delaying the outcome following the response that generated it. For example, the left panel of Figure 7 illustrates the terminal rates of lever pressing by hungry rats obtained by Dickinson and colleagues when each lever press produced a food outcome after a delay of 16, 32, or 64 s (Dickinson *et al.*, 1992). With the 16-s delay and a 20-s memory sample used in our model, only outcomes generated by responses during the first 4 s of a sample occur in the same sample as their responses, whereas with the 32-s and 64-s delays all the outcomes occur in a different sample, thereby reducing the experienced rate correlation. The simulations displayed in the right panel of Figure 7 confirm this intuitive prediction.

**Figure 7.**
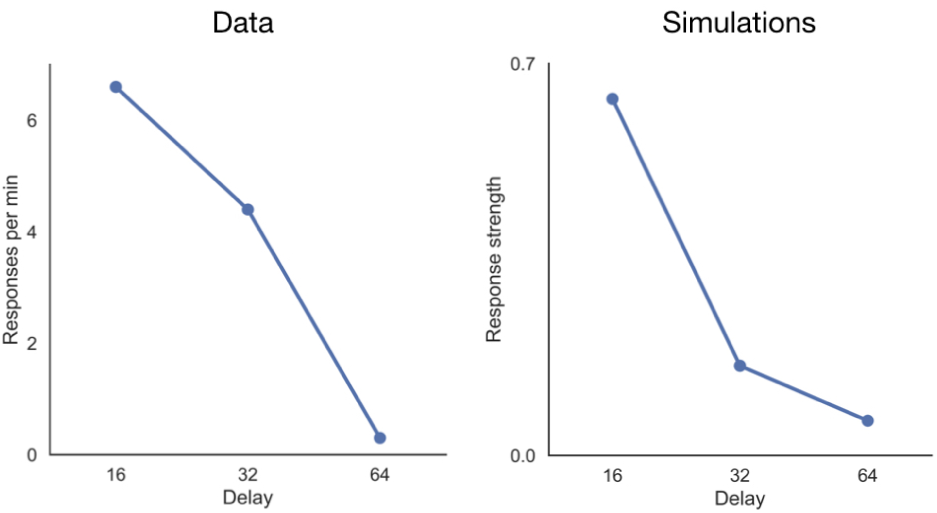
Simulations of rate correlation models for delayed rewards. The left panel shows the data obtained by Dickinson et al. (1992) in rats. The right panel shows simulations of a rate correlation model for the same delay parameters used in the original paper.

Because the response strength *g* also determines goal-directed strength, the model anticipates lower causal beliefs of the response-outcome association when outcomes are delayed. This prediction has been confirmed by Shanks and Dickinson (1991) using fictitious credits as the outcome and key presses as the instrumental response. Moreover, if a causal response-outcome belief underlies goal-directed control, our model anticipates a weaker effect of devaluation on responding when outcomes are delayed. This prediction has been recently confirmed by Urcelay and Jonkman in rats (2019), who found that delaying a food outcome by 20 s after a lever press in one group abolished sensitivity to outcome devaluation compared to a group that underwent training with no delay between the lever response and the outcome.

### Contingency degradation

At first sight, the most direct evidence for a rate correlation approach to instrumental learning is the sensitivity of free-operant performance to the causal response-outcome relationship, in that its computation provides a measure of this perceived contingency. However, the strength of the causal relationship between response and outcome in the environment can be varied not only by changing the probability of a contiguous outcome as in Mazur’s (1983) experiment, but also by varying the likelihood that the outcome will occur in the absence of the action—or the probability of non-contiguous outcomes. When the contiguous and non-contiguous probabilities are the same, the agent has no control over the number of outcomes received in any given time period and its actions are independent to outcome delivery. Hammond (1980) was the first to study the effect of such manipulation in a free-operant procedure. Using rats, Hammond fixed the probability of a contiguous outcome for the first lever press in each second while varying the probability of delivering a non-contiguous outcome at the end of any second without a lever press. Non-contingent schedules, in which the contiguous and non-contiguous outcomes probabilities were the same, failed to sustain lever pressing initially established without the delivery of non-contiguous outcomes.

In general, however, one cannot be certain that a low rate of lever pressing under a non-contingent schedules is due to the absence of a causal relationship between the action and the outcome. Inevitably, the non-contingent schedule greatly increases the frequency of the outcome and therefore the time required to handle and process the outcome with the result that the depression of responding under a non-contingent outcome may have been due to interference with lever pressing by the enhanced outcome handling and processing. One way of addressing this issue is to use a non-contingent schedule while varying the identity of the contingent and non-contingent outcomes. When the contiguous and non-contiguous outcomes are the same, the agent has control over neither the outcome rate nor its identity. However, when the outcomes are different, the agent can control the type of outcomes received because by responding the relative frequency of the contiguous outcome increases.

To illustrate the simulation of contingency degradation by the rate correlation model, we followed an experiment reported by Balleine and Dickinson (1998). Hungry rats were initially trained to lever-press for one of two different food outcomes on an RR 20 schedule so that the probability of the contiguous outcome was 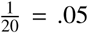. The instrumental contingency was then degraded by delivering a non-contiguous outcome with the same probability of .05 in each second without a lever press. As the left panel of Figure 8 shows, the rats pressed at a higher rate if the non-contiguous and contiguous outcomes were different rather than the same. The right panel illustrates that the rate correlation model can replicate this effect on the assumption that different outcomes receive distinct representations in memory with separate response strengths being calculated for each outcome type.

**Figure 8.**
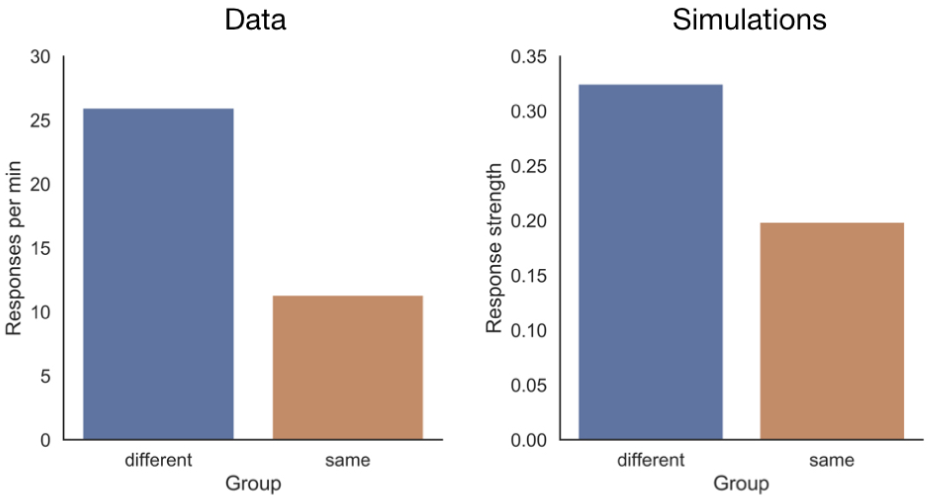
Simulations of a contingency degradation experiment. Left panel. Data obtained by Balleine and Dickinson (1998) in rats. Group *diff* was given freely an alternative outcome with the same probability as the outcome produced by the target action. Group *same* was given freely the same outcome as that produced by the target action. Right panel. Simulations of a rate-correlation model for a similar procedure.

That a causal response-outcome belief is affected by contingency degradation has been demonstrated by numerous studies showing that human causal judgments of the response-outcome association and the rate of responding are lower when the contingency between the response and the outcome is degraded by increasing the probability of non-contiguous outcomes (see Shanks and Dickinson, 1991).

### Interim summary

In summary, this set of simulations demonstrate that our rate-correlation model can in principle provide an account of the primary determinants of instrumental performance: the impact of outcome probability, rate and delay on instrumental performance. In addition, the model correctly anticipates the ratio-interval schedule effect when the outcome probabilities are matched and the effect of degrading the causal contingency between the response and the outcome, both of which are prerequisites for any theory of goal-directed control.

It is equally clear, however, that a further learning system is required for a complete account of instrumental behavior. To the extent that goal-direct learning is assigned to a rate correlation system, we are left with no account of responding on an interval scheduleor ratio schedules with high ratio requirements, both of which are able to sustain responding in spite of establishing very low response-outcome rate correlations. Furthermore, rate-correlation theory on its own provides no principled explanation of why responding extinguishes when outcomes are withheld. As we have noted, the rate correlation cannot be computed at a memory recycle if no outcomes are represented in memory (as would be the case during extinction), and under this circumstance the response strength remains constant at the value computed at the last recycle in which the memory contained at least one outcome representation. A comprehensive account of instrumental action therefore requires an additional learning system.

### A Dual-System Model

When Dickinson (1985) first argued that a rate correlation account of instrumental action could explain goal-directed learning, he embedded it within a dual-system framework to explain instrumental responding that is autonomous of the current value of the outcome, as assessed by the outcome revaluation paradigm. He envisaged this second system as a form of habit learning that involved the acquisition of an association between the stimuli present during training and the instrumental response. This is, of course, the form of stimulus-response learning first postulated by Thorndike (1911) in his original Law of Effect more than a century ago. According to Thorndike, the occurrence of a contiguous attractive outcome following a response simply serves to strengthen or reinforce the stimulus-response association so that the representation of the training stimuli are more likely to elicit the response. However, because all information about the outcome is discarded once it has served its reinforcing function, any subsequent change in the value of the outcome cannot impact on instrumental performance without re-presenting the revalued outcome contingent upon responding. For this reason, to test whether an outcome representation exerts goal-directed control over responding, the outcome devaluation paradigm tests responding in the same training context but in the absence of the now-devalued outcome. Any decrease in responding under these conditions indicates that a representation of the outcome controls an action in accord with the current value of the outcome, thereby demonstrating its goal-directed status (Balleine and Dickinson, 1998; Dickinson and Perez, 2018). Conversely, responding that is impervious to outcome revaluation is generated by a habit system that does not encode the identity of the outcome.

More recent models in the computational RL tradition have attempted to offer a computational account using a similar dual-system approach. RL models recognize two types of systems that closely resemble the two psychological processes just described. Both RL systems aim to maximize the number of rewards obtained by the agent during a task (Daw *et al.*, 2005; Dolan and Dayan, 2013; Keramati *et al.*, 2011). As we have already noted, model-based computations learn a model of the environment by estimating the probability that an action performed in a given state will lead to each of the possible next states and the probability of a reward in each state. As it is this RL system that generates goal-directed action, it is this system that we argue should be replaced by response-outcome learning based on the rate correlation, at least in the case of free-operant behavior.

In addition to a model-based system, RL models recognize another system that is relatively impervious to outcome revaluation. Model-free computations estimate the value of each action in each state (*Q*(*action|state*)) by computing a reward-prediction error in each trial, which is equivalent to storing a running average for the outcome rate obtained by each action in a given state (Sutton and Barto, 1998). Because all the history of rewards is collapsed in *Q*(*action|state*), every time the agent visits a state it maximizes the outcome rate by simply looking up a table with stored *Q*−values and selecting the actions with a higher *Q*−value. For this reason, model-free computations are less computationally expensive and faster than model-based computations (Dezfouli and Balleine, 2013; Keramati *et al.*, 2011). However, when an outcome is revalued or the response-outcome contingencies modified, model-free computations can only adjust to these new conditions by re-experiencing the new conditions of the environment. In this important respect, therefore, the behavioral control exerted by a model-free RL system is similar to the psychological definition of habitual behavior contained in Thorndike’s Law of Effect.

In the following sections, we formalize a dual-system model in which a goal-directed system computes a response-outcome rate-correlation to generate goal-directed strength and interacts with a habit model-free RL algorithm based on a summed reward prediction-error to explain behavioral performance and system control under free-operant training. We show how this model can explain all the empirical phenomena we have already discussed, along with additional results from the literature that are not currently fully captured by RL or associative models of instrumental learning.

As we have already demonstrated a goal-directed system based on the experienced response-outcome rate correlation can capture the primary determinants of free-operant performance. In this section, our goal is to specify a habit algorithm that integrates with this correlation-based goal-directed system to explain behavioral performance on free-operant schedules. To this end, we postulate a model-free algorithm similar to those employed in the RL literature, but modified to account for free-operant procedures. In essence, every time a response is performed, the algorithm deploys a reward prediction-error depending on whether the response was, or was not, reinforced. The value of this prediction-error then changes the habit strength for the next time-step.

Let 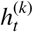 denote habit strength at each time-step *t* in the current memory cycle *k*. In our habit system, the acquisition and extinction of habit strength in cycle *k* follows the following equation:

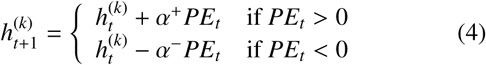

where *α*^+^ and *α*^−^ are parameters between 0 and 1 and represent the learning rates for excitation and inhibition of the stimulus-response connection, respectively, ^4^ and *PE*_*t*_ is the reward prediction-error at time-step *t*, defined as:

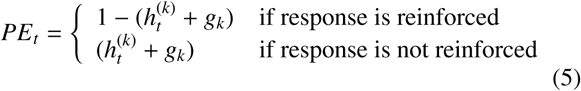

Based on evidence that learning rates for rewarded and non-rewarded episodes are asymmetric (Behrens *et al.*, 2007; Gershman, 2015; Palminteri *et al.*, 2017), we assume that the learning rate of a reinforced response is higher than the learning rate for a non-reinforced response (*α*^+^ > *α*^−^). This assumption is also necessary from a practical perspective: under the partial reinforcement schedules of free-operant schedules, the increment in habit strength produced by reinforcement needs to counteract the effect of a much greater proportion of non-reinforced responses in order to sustain positive levels of responding. Under this habit algorithm, every reinforced episode strengthens the connection between the context and the instrumental response when the reward prediction-error, given by 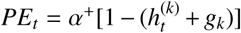 is positive. Likewise, every non-reinforced episode weakens the strength by 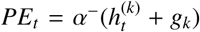. At the end of each cycle, the habit strength is accumulated and used as the starting habit strength value for the following cycle. We denote the habit strength accumulated up to cycle *k* as

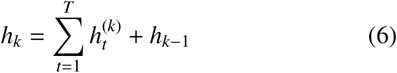

where T is the size of the new sample that is added in each cycle (20 seconds in our particular formulation).

As can be appreciated, the habit system in our model employs a summed prediction term by combining the current response strengths generated by the goal-directed and habit systems. The rationale for this summed prediction term lies with the fact that a prediction-error is intended to capture the extent to which an outcome (or its omission) is surprising or unexpected with respect to the predictions from both systems. In this respect, the rationale for the summed prediction-error in our habit algorithm is the same as the one present in the Rescorla-Wagner rule (1972) for determining associative strength in Pavlovian Learning.

It is important to clarify from a psychological perspective how the interaction between habit and goal-directed systems transfer to instrumental performance in our model. At the end of cycle *k*, and before determining responding for the next cycle *k* + 1, the agent has computed two values: *h*_*k*_ and *g*_*k*_. The activation function takes the linear sum of these values to determine the total response strength for the following cycle, *p*_*k*+1_, which remains constant across this new cycle (i.e., the new time sample added to memory in cycle *k* + 1). However, it is also true that these two computations are not completely independent. First there is the inclusion of goal-directed strength that partially determines the total prediction in the PE term on the habit system. But it is likewise clear that the memory recycling process of the goal-directed system needs to interact with the habit system by constraining the deployment of the accumulated habit strength to the new memory sample that is added after each cycle that elapses. Thus not only there is an independent additive relationship to determine performance, but also an additional interaction during the agents’ behavior within a memory cycle.

Being a model-free algorithm similar to those in the RL field which assign the value *Q*(*action|state*) to a specific action in a given state, Equation 4 assigns response strength to the habit system according to the value of *h*, which completely summarizes the history of reinforcement in a particular state or context. Given that this algorithm is only driven by *PE*_*t*_, it does not explicitly model the information regarding the relationship between the response and the outcome or its current motivational value, making it insensitive to manipulations of outcome value or the causal relationship between the response and the outcome. Such behavioral autonomy is the cardinal feature of habitual behavior (Dickinson, 1985).

Having specified the algorithm for the habit system, the next step is to specify the type of interaction between behavioral systems that would explain performance and behavioral control for different experimental conditions in free-operant training. Our assumption in this regard will be motivated by the data reported by Dickinson et al. (1983, see Figure 1). After training two groups of rats under interval and ratio schedules with matched outcome probabilities, Dickinson and colleagues devalued the outcome in half of the rats in each group by pairing it with toxicosis. After the devaluation manipulation, only the ratio-trained rats decreased responding (i.e., were under goal-directed control). By contrast, the performance of the interval-trained rats at test was unaffected by outcome devaluation (i.e. they were under habitual control). An interesting feature of these data is that the level of responding after devaluation in the ratio-trained group did not differ from that of the interval-trained group. Because the outcome probability was matched between the groups, the habit system’s contribution to responding should have been equal in both groups. Likewise, because by definition responding that is sensitive to devaluation must be attributed to the goal-directed component, the residual responding that was not affected by devaluation in the ratio-trained group must, by necessity, be attributed to a habitual component that was still active after devaluation of the outcome. In light of these data, we propose a model where the relative strengths of each system concurrently contribute to response probability, which we denote by *p*. In particular, we assume that response probability in cycle *k* + 1, *p*_*k*+1_, is governed by a sigmoid function:

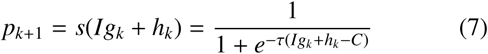

where *g*_*k*_ is the goal-directed strength in cycle *k* as defined above, *h*_*k*_ is the habit strength accumulated by Equation 4 during the experiment, up to cycle *k*, and *I* is a variable representing the current incentive value of the outcome by taking the value 1 if the outcome is valued and 0 if the outcome is devalued (that is, we assume that the devaluation procedure successfully decreases the incentive value of the outcome to zero). The parameter τ is an inverse temperature parameter that reflects how sensitive the agent is to increases in total response strength (*Ig*_*k*_ + *h*_*k*_)) and *C* is a bias parameter (*C* = 0.6) which represents a bias towards responding over waiting. Under this total response function, the two systems summate to determine total responding, and therefore response probability in the next cycle *p*_*k*+1_ reflects the relative contribution of each system to behavior (see Figure 9).

**Figure 9.**
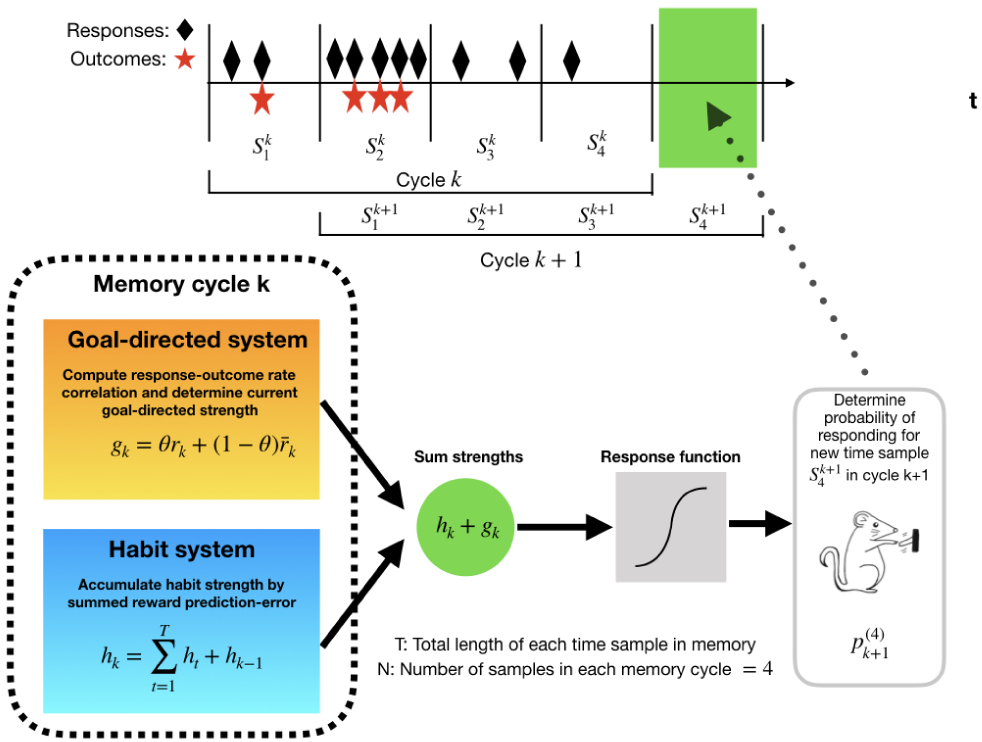
Schematic representation of the dual-system model. For each cycle, the agents concurrently computes the response-outcome rate correlation while habit strength is accumulated. The strength of both systems is then summed and a response activation function produces the probability of responding for the new sample in the following cycle. The rate correlation on the goal-directed system is only computed when both responses and outcomes are held in memory ; otherwise, the system is inactive and the value stays constant at the last value registered in memory. (Illustrations courtesy of Loreto Contreras.)

In the following sections, we will discuss the implications of such an assumption for the type of behavioral control that should be expected for different free-operant procedures by performing simulations of this dual-system model. We should note that the aim of these simulations was not to fit the free parameters of our model to the experimental data presented here. Instead, we performed an unsystematic search for those parameters that predicted the qualitative patterns of the data that originally motivated our theory (i.e., Dickinson *et al.*, 1983) and kept the values of those parameters constant across all simulations (see Niv *et al.*, 2005; Soto *et al.*, 2014; Soto and Wasserman, 2010; Wagner, 1981 for similar approaches).

### Ratio and interval training

Initially, we simulated goal-directed and habitual learning under interval and ratio contingencies using a RI 15-s master schedule. The outcome probability generated by each master interval simulation was then used to generate a yoked simulation on a ratio schedule with a parameter that yielded the same outcome probability. The initial response probability for the first session of training reflected one session of pretraining under RR 5 and each session was terminated after 30 outcomes had been received.The response rate for the first memory cycle was set to 10 per min. Panels a and b of Figure 10 display the mean values generated by 200 simulations under the master interval and yoked ratio schedules, respectively. Shown separately are the response strengths generated by the goal-directed and habit systems, *g* and *h* respectively, and the resultant probability of responding per second, *p*, produced by the cooperative interaction of these response strengths.

**Figure 10.**
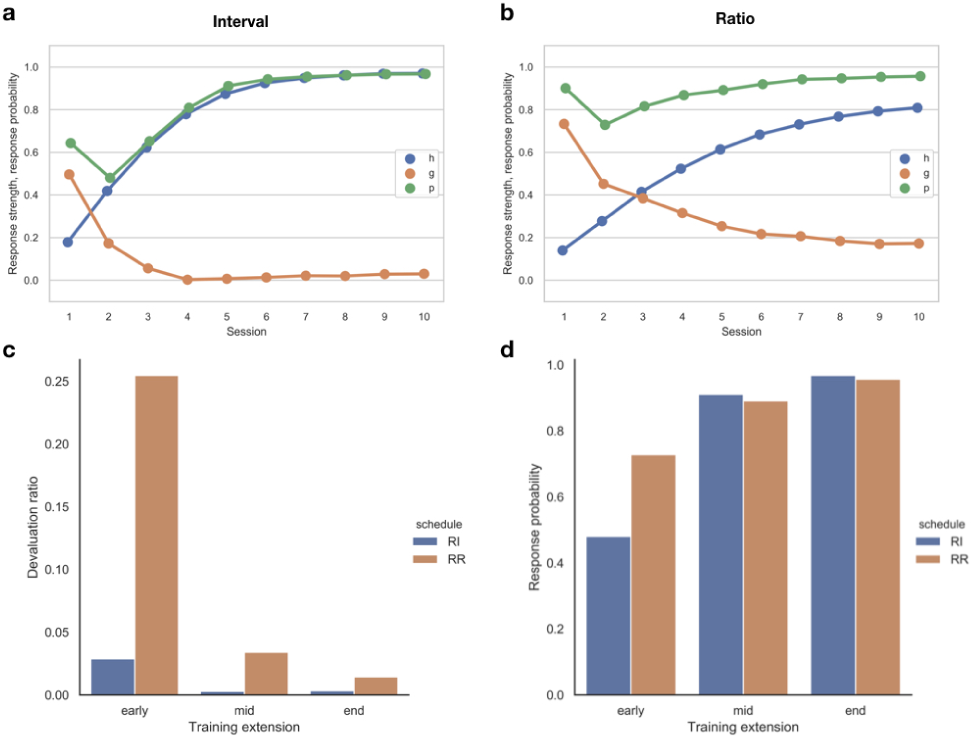
Simulations of the dual-system model for the experiment reported by Dickinson et al. (1983). (a) Strength from each system and response probability per second across 10 sessions of interval training. (b) Strength from each system and response probability per second across 10 sessions of training under yoked ratio training, matching outcome probabilities with the interval-trained subjects. (c) Sensitivity to outcome devaluation for ratio and interval training as assessed by a devaluation ratio early in training (Session 2); at mid-training (Session 5) and at the end of training (Session 10). (d) Response probability per second for ratio and interval training across different extensions of training.

The first point to note is that the model reproduces the differential sensitivity of ratio and interval performance to outcome devaluation early in training. For example, ratio training generates a goal-directed response strength, *g*, of about 0.4 by the third session, whereas the interval response strength is close to zero for equivalent training. As the model assumes that outcome devaluation, if complete, abolishes the contribution of *g* to overall responding by setting *I* = 0, the model naturally explains why devaluation has a greater impact on ratio than on interval responding early in training (Dickinson *et al.*, 1983). This finding is summarized in Figure 10c in terms of a devaluation ratio, *DR*, defined as 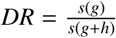, where *s* is the sigmoid function as defined in Equation 7.

### The development of behavioral autonomy

Perhaps, however, the most notable feature of these simulations is the decline in the goal-directed response strength as the habit strength grows with training. This reduction in *g* reflects, at least in part, the reduction in the variance of the rate of responding across the time samples in memory: as the overall response rate increases, the experienced rate correlation, and therefore *g*, declines. Thus, according to our model, behavioral autonomy should develop as responding becomes stereotyped with more extended training. The reduction in sensitivity to outcome devaluation with training predicted by the simulations is documented in Figure 10c in terms of a devaluation ratio calculated for three different training extensions.

Adams (1982) was the first to report that behavioral autonomy (i.e., habitual behavior) developed with training on a variety of ratio schedules. Although the development of autonomy with training has been independently replicated multiple times (e.g., Dickinson *et al.*, 1995; Holland, 2004; Killcross and Coutureau, 2003), a number of studies have reported goal-directed control even after extended training. For example, de Wit et al. (2018) have documented two failures to replicate the development of behavioral autonomy observed by Tricomi et al. (2009) after training humans under an interval schedule (see Corbit *et al.*, 2014; Nelson and Killcross, 2006). Similarly, Jonkman et al. (2010) found that rats remained sensitive to outcome devaluation throughout 20 sessions of interval training and Garr et al. (2019) have recently reported an outcome devaluation effect that only emerged after extended training on an RI schedule. We shall return to these problematic effects when we discuss motivational effects on free-operant responding.

### Choice training

Our analysis of extended training makes clear that, according to rate correlational theory, the conditions for developing habitual control are not directly determined by the reinforcement schedule nor the amount of training but rather by whether or not the agent experiences a correlation between the rates of responding and outcomes as represented within the memory cycle. To recap, embedding rate correlational theory within a dual-system model predicts a reduction in the experienced rate correlation through the development of invariant stereotyped responding with the growth of habit strength, an effect enhanced in the case of interval schedules by the temporal control of outcome availability.

The cardinal importance of the experienced rate correlation is reinforced by the contrast between the single-response training, which has been our focus so far, and free-operant choice or concurrent training, which involves interleaved experience with two different response-outcome contingencies (Colwill and Rescorla, 1985; Colwill and Rescorla, 1988). However, of more direct relevance to the present analysis is the study by Kosaki and Dickinson (2010), which we have already discussed briefly with respect to the differential reinforcement of long IRTs, in that they directly compared behavioral autonomy after concurrent and single-response training.

To recap, Kosaki and Dickinson (2010) trained rats on two RI schedules that were concurrently active during each session of training. In one group, the *choice* group, responding on different levers produced different outcomes. Another group of rats, the *single-response* group, received the same two outcomes, except that in this group one of the outcomes was earned by responding on one lever, whereas the other outcome was delivered after the same average period of time but independently of responding (i.e., non-contingently). After 20 sessions, a contingent reward was devalued in both groups by aversion conditioning and responding tested in a subsequent extinction session. Kosaki and Dickinson (2010) observed that responding in the single-response group was insensitive to devaluation, whereas the choice group markedly reduced the rate of the response whose outcome was devalued.

There are two points to note about Kosaki and Dickinson’s (2010) findings. First, the devaluation effect was assessed against control conditions in which the other outcome was devalued. As a consequence, any effect of outcome devaluation that was not mediated by the instrumental contingency, such as the motivational effect of contextual conditioning on general performance (see below), was equated across conditions. Second, the same devaluation effect was found whether or not the choice was tested with both levers present or just a single lever. Thus, the devaluation effect exhibited by the choice group arose from the training rather testing conditions. In conclusion, these results demonstrated that responding in the choice group was still under goal-directed control even when similar extended training rendered responding habitual in the single-response group.

Recall that, according to the rate correlation component of our dual-system model, habitual behavior develops through extended training because responding becomes stereotyped with little variation across time-samples, thereby yielding a low rate correlation experienced within a memory cycle, an effect compounded by the intrinsic low rate correlation engendered by an interval contingency. However, response rate variation across time-samples is an inevitable consequence when the agent is engaged with two interval sources of reward. When engaged with one of the sources, the memory samples will register neither responses nor outcomes from the non-engaged source. Consequently, any memory cycle containing a switch will have some samples with no response nor outcome representations of the switched-to-source and other samples containing these representations in the active source. Of course, the same will be true of the switched-from-source. As a consequence, the agent will experience a sustained rate correlation for both responses, each of which will sustain goal directed control. To substantiate this intuitive analysis, we simulated a concurrent choice procedure similar to that employed by Kosaki and Dickinson (2010). The simulations were run under the same conditions as the previous ones for interval training in the previous simulation of Dickinson et al.’s (1983) experiment; that is, the initial response rate was set to 10 per min, followed by one session of pre-training under RR 5.

It is well established that the probability of switching away from a source remains constant during responding to that source (Heyman, 1979) and so we programmed a fixed probability per 1-s time sample, *p*_*switch*_, for a change-over between levers in the case of the *choice* group. Inspection of the authors’ original data-set revealed that their rats switched between levers on average every 10 s; we therefore set *p*_*switch*_ = .1 for the following simulations.

Figure 11 shows the results of the simulation by the dual-system model for this choice experiment. As can be seen, similar amounts of training under a choice procedure yield significant contributions of the goal-directed system compared to single training. The result holds even when the amount of training is sufficient to drive the habit strength to asymptote, a factor that should reduce the experienced rate correlation (and hence goal directed control) if only a single response was available. In summary, the model predicts that both systems should contribute to the control of responding under choice training, and therefore outcome devaluation should be effective in modulating responding under choice procedures, in line with the results reported by Kosaki and Dickinson (2010).

**Figure 11.**
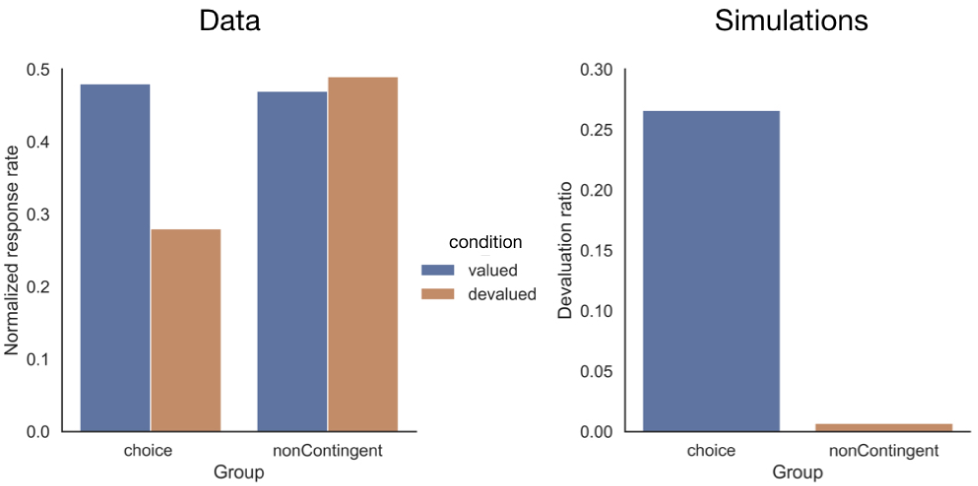
Simulations of Kosaki and Dickinson (2010), investigating sensitivity to outcome devaluation in a choice procedure. The left panel shows the results obtained by the authors and the right panel the results of the simulations of the dual-system model. The choice group was trained concurrently with two responses yielding two different outcomes under RI schedules with the same interval parameters.

One thing to note with regard to Kosaki & Dickinson’s (2010) study is that the outcomes produced by each response differed in their sensory properties, which is critical if the dual-system model is to predict devaluation sensitivity after overtraining. It is an important feature of the goal-directed system that separate rate correlations are calculated for each type of outcome experienced. Using the same outcome for each of the responses effectively changes the schedule into a non-contingent one for both responses because the outcome rate when the agent is responding to one source would be the same as that when response are not directed at that source. Hence, the rate correlation for this response should be close to zero with the consequence that responding under such a schedule should be purely habitual. Holland (2004, Experiment 2) conducted an experiment where the same training regime was given to two different groups of rats under interval schedules, with two different responses and outcomes available in one group, and with two responses producing the same outcome in another group. After extended training, only the rats in the group trained with multiple outcome was sensitive to devaluation; using a single outcome even when two responses were available made responding autonomous of the current value of the outcome, in line with the predictions of our model.

### Temporal scope

Although our dual-system model provides a plausible account of the Kosaki and Dickinson (2010) demonstration of the persistence of goal-directed control under extended training choice or concurrent training, Colwill and Rescorla (1985) also reported such persistence under training conditions that are more problematic for our account. Importantly, they trained the two response-outcome interval relationships in separate sessions before giving their rats a choice test in extinction with both response manipulanda present following the devaluation of one of the outcomes. The reason why this finding is problematic for our implementation of rate correlation theory in a short-term memory system is because exposure to the conjoint absence of a response and its outcome in sessions when the other response-other contingency is being trained is not sufficiently temporally contiguous to the sessions in which this response is trained with its outcome. As a consequence, the conjoint absence of the response and its outcome and their conjoint presence cannot be plausible represented within the same short-term memory cycle, a condition that is necessary for a sustained positive rate correlation for that response and its outcome.

A similar problem for our short-term memory system for assessing the local experienced rate correlation arises from Adam’s (1982) initial analysis of behavioral autonomy. Having initially established lever-pressing by all his rats (Adams, 1982, Experiment 5—see also Dickinson *et al.*, 1995), he then overtrained one group on a variable ratio schedule, which yielded habitual behavior. By contrast, the experimental non-contingent group received the same number of outcomes but the absence of the lever except for the last session when lever pressing was trained on the variable ratio schedule. Like the overtrained group these animals showed habitual behavior even though the amount of instrumental training they received was such as to render responding by a control group sensitive to outcome devaluation in the absence of the prior non-contingent exposure. Once again, it appears that the animals are capable of integrating information about the outcome from sessions without the opportunity to perform the target response with that gained in sessions with the opportunity to respond in a way that modulates the experienced rate correlation appropriately. In the case of Colwill and Rescorla (1985), interleaved concurrent training exposure to training context in the absence of the outcome enhances the experienced rate correlation whereas context exposure with non-contingent outcomes reduces the experienced rate correlation in Adams’ (1982) experiment.

Although we do not have the empirical evidence to specify the psychological mechanisms by which such integration takes place, we may assume that the mechanism is functionally equivalent to increasing the size of the memory across which the experienced rate correlation is assessed. For example, it is possible that the return to a context in which the target response has been previously trained leads to the retrieval of the contents of the final memory cycle in the same context during the previous session, which is then integrated with contents of the the initial memory cycle established by training the target response. To illustrate how such a system may operate, we performed simulations of the dual system model with an enhanced memory on a non-contingent pre-exposure schedule similar to that employed by Adams (1982).

The left panel of Figure 12 presents the results obtained by Adams expressed as a devaluation ratio. To simulate this experiment, we increased the number of time samples deployed by subjects to 500 samples. For the first cycle of 500 samples, we delivered free outcomes with *p*_*f ree*_ = .2, which is equivalent to an RT 5 sec schedule; a single session of RR 10 training then followed. The session was terminated after 30 outcomes were delivered. For the control group, we simulated the first cycle of 500 memory samples with a probability of responding *p*_*initial*_ = .1, followed by a single session of RR 10 training. The results of these simulations are shown in the right panel of Figure 12. Although using a larger memory size did not abolish the devaluation effect completely, as observed by Adams (1982), the devaluation ratio for the *pre-trained* group is lower than that for the *control* group, anticipating a weaker devaluation effect after outcome pre-training.

**Figure 12.**
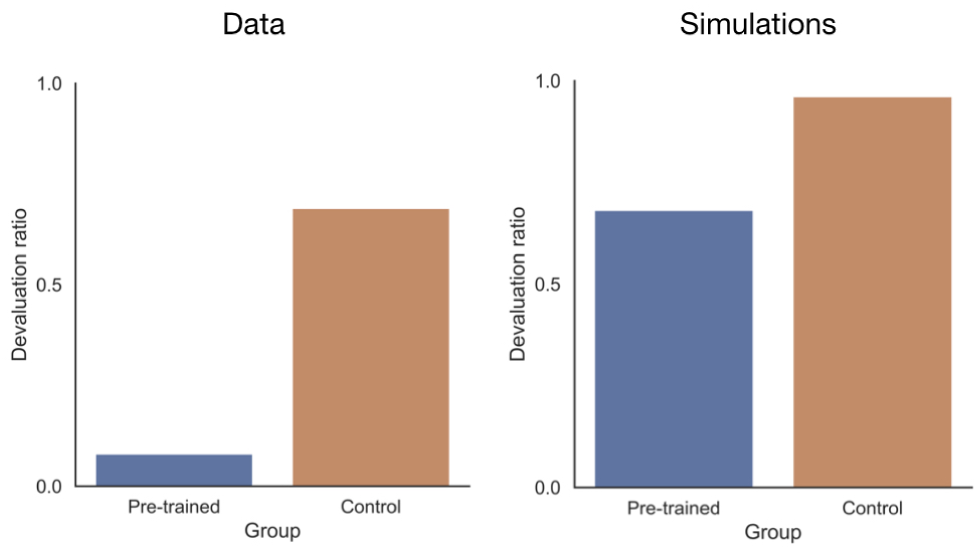
Simulations of an experiment run by Adams (1982, Experiment 5) where one group was given extended magazine training before a single session of training under RR 10. The left panel shows the results obtained by Adams during the extinction test, measured by a devaluation ratio. The right panel shows the simulations of the dual-system model for a similar procedure, employing and increased memory size.

### Extinction

As it stands, the rate correlation system in our model makes what at first sight appears to be a highly problematic prediction: goal-direct control should never extinguish. Recall that the goal-directed system only computes the response-outcome rate correlation for memory cycles in which at least one response and one outcome are registered in memory. The consequence of this assumption is that goal-directed strength remains frozen throughout extinction at the the level attained during acquisition following the last memory cycle that contained an outcome representation.

Although not generally acknowledged by RL theory, this prediction accords with a series of studies conducted by Rescorla (1993), who reported that the impact of the outcome devaluation is not reduced by extinction. In one of his experiments, Rescorla trained two responses each with a different outcome, and then one of the responses was extinguished before a final devaluation test. Having re-established responding with a third outcome, Rescorla found that devaluating one of the original training rewards produced a comparable reduction in performance of the associated response in extinguished and non-extinguished conditions, thereby demonstrating that goal-directed learning survived the extinction phase. The left panel of Figure 13 presents the comparable outcome devaluation effect observed by Rescorla (1993) in the extinguished and non-extinguished conditions.

**Figure 13.**
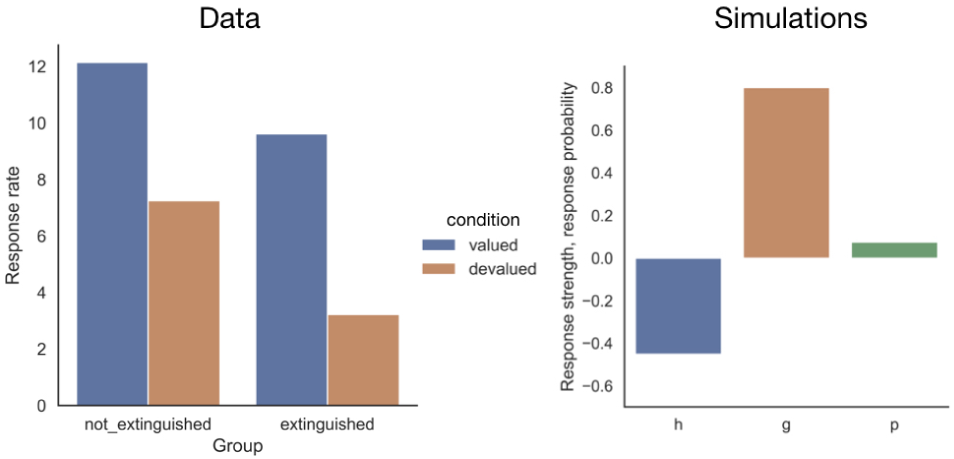
Simulations of a devaluation manipulation after extinction. The left panel shows the results reported by Rescorla (1993), which involved devaluation of one outcome for one response after an extinction phase compared with a response for which the outcome was not devalued. A control group had similar training but without undergoing an extinction phase. Note that before the devaluation test, responding was recovered by training with another outcome after extinction. The right panel shows the final values after one session of RR 5 followed by 2000 cycles of extinction for the dual-system model.

The most plausible explanation for this result was offered by Colwill (1991), who reasoned that the goal-directed strength remains unaffected during the extinction phase because it is inhibited by the habit system when outcomes are withheld. Our dual-system model anticipates the acquisition of this inhibitory habit strength during extinction following a similar reasoning. In our model, the prediction error term in the habit system includes the total prediction determined by the linear sum of the habit and goal-directed strengths *g* and *h*, respectively. Assume that cycle *k* is the last one containing response and outcome events in memory (the last cycle in training, in this example). If *g* retains a positive value during extinction (because the response-outcome rate correlation is not computed in a memory cycle that does not contain any outcomes) then *g*_*k*_, = *g*_*k*_ = *g*_0_ for all cycles *k*, in extinction. Then it follows that the prediction-error for *h* will be negative at each time step (*PE*_*t*_ =− (*h*_*t*_ + *g*_0_)) and hence there will be a systematic decrease of *h* during extinction. The reductions in *h* will in turn decrease *p* with training, and responding will eventually extinguish. Indeed, and in accord with Colwill’s hypothesis, the habit strength *h* will have to become negative or inhibitory for complete extinction to occur, because *g* remains constant and positive throughout the extinction phase.

To simulate extinction, we initially trained our virtual rats as in the pre-training stage of our previous simulations; that is, the response rate for the initial memory cycle was set to 10 per min followed by a single session of training under RR 5. Outcome delivery was then suspended for 2000 additional memory cycles. As can be appreciated in the right panel of Figure 13, the model correctly predicts a systematic decrease in total responding while maintaining a positive goal-directed strength, providing an account of the retention of goal-directed control reported by Rescorla ^5^.

The survival of *g* across extinction raises the question as to which training conditions would have an impact in goal-directed responding in our model. According to our account, goal-directed responding can only be extinguished by exposure to a non-contingent schedule, so that the response-outcome rate correlation experienced by the agent is degraded (see Figure 8). There is evidence that exposure to non-contingent outcomes does not reduce outcome-specific Pavlovian-instrumental transfer (Colwill, 2001; Rescorla, 1992), but this form of transfer is thought to be unaffected by outcome devaluation (see Cartoni *et al.*, 2016; Holmes *et al.*, 2010). Moreover, transfer learning differs from that mediating goal-directed behavior (1994). As far as we know, the impact of outcome revaluation following a reduction in responding induced by non-contingent training has not been reported and so represents a novel prediction of our model. We simulated this novel prediction by training our agents as in the previous extinction simulations, but switched them from RR 5 to a non-contingent schedule in which free reinforcers were delivered with the same probability as that programmed by the RR schedule (.2). The simulations confirmed our prediction (see Figure 14): degrading the response-outcome contingency significantly reduced the contribution of the goal-directed system that would otherwise stay constant by inhibition from the habit system in an extinction procedure.

**Figure 14.**
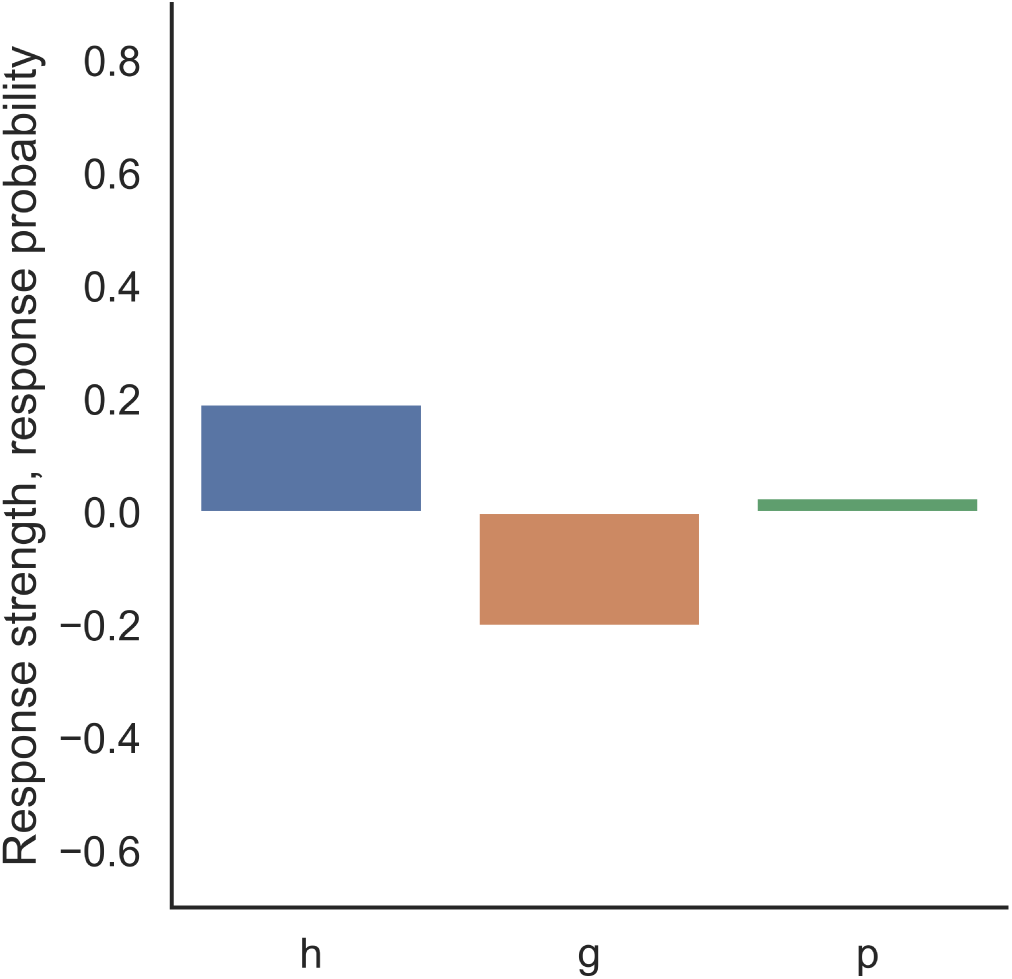
Simulations of a dual-system model for a devaluation manipulation after non-contingent training. The plot shows the final outputs of the systems after one session of RR 5 training followed by 2000 cycles of non-contingent training for the dual-system model.

As can be appreciated from the set of simulations we have presented, our dual-system model can explain a wide range of phenomena from the instrumental free-operant literature. However, there still remain a number of behavioral processes that are not as yet open to simulation by our formal model, but that are in principle compatible with it. We discuss these processes in turn below.

### Performance after extended training

As Figure 10d clearly illustrates, the dual-system model predicts that a ratio schedule maintains a higher response rate than a comparable interval schedule early in training, as was originally observed by Dickinson et al. (1983). With more extended training, however, the difference in performance disappears as responding comes under habitual control. This prediction is clearly at variance with the empirical evidence showing a sustained ratio-interval performance effect with extended training (e.g. Catania *et al.*, 1977; Pérez *et al.*, 2016; Pérez and Soto, 2020). We have already noted that interval schedules differentially reinforce long IRTs—the longer an agent waits before responding again, the more likely it is that a further outcome has become available with the resultant increase in the probability of reinforcement (see Figure 2a). To the extent that habit learning is conceived of as a form of stimulus-response learning, we should expect this form of learning to be sensitive to the temporal cues registering the time since the last response and hence to come under the control of these cues, with the resulting impact on the rate of responding. By contrast, on a ratio schedule the probability of reinforcement is constant and independent of IRTs, so responding should be independent of the size of the emitted IRTs (for a discussion, see Chapter 4 in Mackintosh, 1974).

As the habit system does not incorporate a mechanism for the differential stimulus control of responding, we cannot use our model to assess impact of IRT reinforcement on responding. However, if this differential reinforcement could be removed while implementing the low rate correlation characteristic of interval contingencies, the model predicts that there should be no sustained ratio-interval performance effect. Kuch and Platt (1976) specified such a schedule, now referred to as a regulated-probability interval schedule (RPI). Without going into the implementation details, the RPI schedule sets the probability of reinforcement for the next response so that if the agent continues responding at the current rate, the rate of the outcome will match that specified by the scheduled interval parameter. As a consequence, variations in the rate of responding will have little impact on the obtained outcome rate. In other words, the schedule maintains the low rate correlation characteristic of a standard interval schedule. However, as the outcome probability for the next response is fixed at the time of the preceding response, the RPI schedule, like a standard RR schedule, does not differentially reinforce any particular IRT. Consequently, our dual system model predicts that there should be no difference in the sustained responding on ratio and RPI schedules with matched outcome rates or probability.

The limited empirical evidence on this contrast is mixed. Neither Tanno and Sakagami (2008) nor Perez et al. (2018), who both trained hungry rats to lever-press for a food outcome, reported a sustained difference between responding on ratio and matched RPI schedules, while observing the reduced response on a standard matched interval schedule. In contrast, Dawson and Dickinson (1990) observed a sustained higher response rate of chain pulling on a ratio schedule than on a yoked RPI schedule. The source of this empirical anomaly is as yet unclear, although it may well be that the use of an atypical chain-pulling response by Dickinson and Dawson produced a sustained variation in the rate of responding relative the prototypical lever pressing. If so, the chain-pulling rats would have continued to experience a higher rate correlation under the RR schedule than under the matched RPI schedule and therefore, according to the model, should have continued to respond at a higher rate due to the sustained goal-direct response strength *g*.

### Motivational processes

Different processes are involved in the motivation of habits and goal-directed action and so we shall consider each in turn in relationship to our dual-system model.

#### Incentive learning

In Equation 6 We have already included the incentive value *I* of outcomes in the term for the contribution of the goal-directed system to determining the response probability in a memory cycle. In the case of biological outcomes, such as foods and fluids, animals have to learn about their incentive values through consummatory experience with these commodities if they are to function as goals of an instrumental action, a process that Dickinson and Balleine (1994; 2002) refer to as incentive learning. Moreover, the animals also have to learn how these incentive values vary with motivational states, such as hunger and thirst. Dickinson and Dawson (1988; 1989) first reported the role of incentive learning in the motivational control of goal-directed action in an irrelevant incentive procedure using a shift from training under hunger to testing under thirst. Their rats were initially trained to lever-press and chain-pull, one for food pellets and the other for sugar water, while hungry. During a subsequent extinction test, thirsty rats only preferentially performed the action trained with the sugar water if they had previously had the opportunity to drink the sugar water while thirsty, indicating that they had to learn about the incentive value of the sugar water when thirsty. Such incentive learning is required not only for shifts between motivational states but also variations within a motivational state, such as that between satiety and hunger (Balleine, 1992). Dickinson and Balleine (2010) have subsequently argued that the assignment of incentive value to an outcome is based on the experienced hedonic reactions to, and evaluation of that outcome.

#### Motivating habits

In accord with classic two-process theory (Rescorla, 1967), it is well established that Pavlovian stimuli associated with appetitive reinforcers motivate free-operant behavior established and maintained by appetitive outcomes. Estes (1948) was the first to demonstrate this effect using what has come to be called the Pavlovian-instrumental transfer (PIT) paradigm (see Cartoni *et al.*, 2016; Holmes *et al.*, 2010 for reviews). He initially established a Pavlovian stimulus as a signal for food before training his hungry rats to press a lever for the food. When he then presented the stimulus for the first time while the rats were lever-pressing, he observed an increase in response rate during the stimulus. Given this transfer, two-process theory assumes that the Pavlovian conditioning to contextual cues occurs concurrently with instrumental learning during standard operant training so that the context comes to exert a motivational influence on free-operant performance.

The concordance between the impact of outcome rate on operant performance and Pavlovian responding accords with this two-process theory of instrumental motivation. It has long been recognized that an important variable in determining the rate of responding on interval schedules is the outcome rate rather than the outcome probability per response, and Killeen (1978; 1982) proposed that outcome rate has a direct motivational impact so that higher outcome rates will have a general and sustained energizing effect on behavior. Indeed, this effect has been formalized by Herrnstein and collegues (de Villiers and Herrnstein, 1976) in terms of a hyperbolic function between response and reinforcement rates and, more recently, Harris and Carpenter (2011) have reported that the same function applies to Pavlovian conditioning of magazine approach in rats, consistent with the idea that the sensitivity of instrumental responding to outcome rate reflects a motivational influence of Pavlovian contextual conditioning on response vigor.

This Pavlovian motivation modulates habitual rather than goal-directed behaviour. Holland (2004) reported a stronger PIT effect when habitual control had been induced by extended training, whereas Wiltgen et al. (2012) reported a similar association between the habitual status of responding and general PIT in mice by contrasting ratio and interval training. They observed greater PIT following interval training when performance was impervious to outcome devaluation than following ratio training when responding was still under goal-directed control. Further evidence that the target of Pavlovian motivation is habitual comes from the fact that the magnitude of PIT was unaffected by whether the outcome associated with the Pavlovian stimulus was the same as or different from the instrumental outcome ^6^.

A compelling demonstration of the generality of Pavlovian motivation comes from an irrelevant incentive study of PIT. Dickinson and Dawson (1987) trained hungry rats to lever-press for food pellet while also giving separate Pavlovian training by pairing one stimulus with the pellets and another with sugar water in the absence of the lever. When for the first time the rats were given the opportunity to press the lever during the stimuli while thirsty and in the absence of any outcomes, they did so more during the sugar-water stimulus than during the pellet signal. This finding establishes two important points. The first is the generality of the motivational influence which augments any prepotent habitual response even if that response was trained with a reinforcer that differs from that associated with the stimulus. Second, the Pavlovian motivational process can endow habitual responding with a veneer of goal-directedness. The shift of motivational state from training under hunger to PIT testing under thirst is an apparent outcome revaluation procedure in that the sugar-water reinforcer remained relevant to the test motivational state whereas the pellet reinforcer did not. However, this apparent outcome revaluation effect did not indicate goal-directed control because the revaluation did not operate through a representation of the response-outcome contingency in that lever pressing was trained with the food pellets, not the sugar water. Moreover, in contrast to the instrumental incentive learning, it was not necessary for the animals to re-experience the outcome in the new motivational state for a PIT effect to occur: the sugar-water stimulus enhanced lever pressing in the extinction test even though their rats had never previously experienced the sugar water while thirsty.

The sensitivity of this Pavlovian motivation to an outcome revaluation procedure can easily lead to the erroneous attribution of goal-directed status. For example, Jonkman et al. (2010) reported that the rate of lever pressing remained sensitive to outcome revaluation even after extensive training on an interval schedule. It is very likely, however, that the apparent devaluation effect was mediated by Pavlovian contextual motivation of habitual responding. Extinguishing context conditioning prior to the devaluation test significantly reduced the magnitude of the effect. This same Pavlovian process may have operated in two more recent outcome devaluation effects that are problematic for our dual-system model. Recently, Garr et al. (2019) have reported that lever-pressing by rats established with an RI schedule was impervious to outcome devaluation after minimal and intermediate training but became sensitive to outcome value after more extended training. This finding is compatible with the view that the influence of contextual Pavlovian motivation increases with habit strength, so that changes in motivational state have a stronger impact on responding with extended training. Conversely, the lack of devaluation sensitivity after limited training may have been a consequence of weak contextual conditioning.

In the case of the second problematic effect, Coutureau and Killcross (2003) gave extensive lever-press training that was sufficient to establish habitual responding in a subsequent outcome devaluation test for their control rats. By contrast, their experimental animals showed a reliable devaluation effect when the infralimbic cortex was inactivated by muscimol immediately prior to the devaluation test. They interpreted this result as evidence that disrupting the habitual system revealed an underlying goal-directed control. This interpretation is at variance with our model, which argues that behavioral autonomy results from a loss of goal-directed strength through the development of stereotyped invariant responding so that there should be minimal goal-directed control to be reinstated. Coutureau and Killcross (2003) were aware that a spurious devaluation effect could arise through a contextual Pavlovian process and, consequently, their rats received exposure to the operant context without the lever or outcomes to extinguish the contextual conditioning. However, it is possible that the administration of the muscimol may have renewed the contextual conditioning and thereby reinstated the Pavlovian outcome devaluation effect. It is well established that after Pavlovian conditioning and extinction in one context, a shift to another context can renew the Pavlovian response (Bouton and Ricker, 1994). So it may well be that administration of the muscimol acted as a context shift by changing the internal state of the rats with the result that the muscimol, rather than revealing latent goal-directed control, functioned to renew the Pavlovian outcome devaluation effect.

Finally, it is possible that two more recent demonstrations of the reinstatement of an outcome devaluation effect may also be mediated by the motivational impact of Pavlovian contextual conditioning on habitual responding. In one set of studies (Bouton *et al.*, 2020) the reinstatement was induced by presenting a novel food at the end of of instrumental training, whereas in the second set (Trask *et al.*, 2020) interleaved training of another response in a second contextual during the final sessions of training of the target response produced the reinstatement. In neither case, however, was it demonstrated that the reinstated outcome devaluation effect was mediated by the instrumental contingency between the target response and its outcome rather than by Pavlovian conditioning to the context in which the target response was tested. Therefore, we cannot be certain that the outcome devaluation effects reflected a reinstatement of goal-directed control.

As we have noted, the performance function in our dual-system model (see Equation 7), which transforms response strengths into response probability, includes a parameter *I* that represents the current incentive value of the outcome so that *Ig* determines the contribution of the goal-directed system to performance. By analogy, we extend it by also including an additional parameter that reflects the motivational effects of appetitive Pavlovian stimuli on habitual performance. Following Hull’s (1943) classic nomenclature, we denote this parameter as *D* for drive, which multiplies the habit strength *h* to represent the contribution of the habit system to overall performance. Like the Hullian drive concept, *D* appears to exert a general motivational effect, at least within the appetitive domain, so that the complete response function has the form

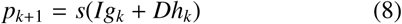

where *s* is the sigmoid function as shown in Equation 7.

In summary, the motivation of habits and goal-directed actions is varied and complex, even in the case of basic biological commodities such food and fluids. Habits are motivated by a general appetitive drive *D* conditioned to contextual and eliciting stimuli, whereas the incentive value of the outcome *I*, which is learned, motivates goal-directed action. Habitual motivation is directly sensitive to shifts in motivation state, whereas the agent has to learn about incentive values of outcomes in different motivational states before they can control goal-directed action.

### Discriminative control

As it stands, our dual-system model offers no mechanism by which goal-directed responding can come under stimulus control as the goal-directed strength, *g*, is solely a product of the experienced correlation between response and outcome rates and the current incentive value of the outcome *I*. There is, however, extensive evidence that such responding can come under discriminative control. The most compelling comes from an elegant biconditional discrimination studied by Colwill and Rescorla (1991). They trained rats with two different responses (R) and outcomes (O) and arranged for the different stimuli (S) to signal which outcome would be produced by each response. When S1 was present, R1 led to O1 and R2 to O2, whereas the opposite relation held when S2 was present (R1 led to O2 and R2 led to O1). When one of the outcomes was then devalued, rats responded more in the extinction test during the stimulus that during training signalled the non-devalued outcome for the target response. As this design equates the stimulus-outcome associations across stimuli and the response-outcome association across responses, this devaluation effect requires the encoding of the triadic relationship between stimulus, response and outcome, a representation that is not as yet incorporated into our current formulation of the model. The nature of this representation has been a long standing issue (Rescorla, 1990). One possibility is that the stimulus is associated with the rate correlation experienced in its presence and thereby modulates the strength of the response-outcome relationship controlling goal-directed responding during the stimulus. As a consequence, the same response can enter into different rate correlations, each of which would be associated with relevant discriminative stimuli. An alternative is that a response is represented in memory in a configuration with the discriminative stimulus and that different rate correlations are computed for each stimulus-response configuration.

Whatever the merits of these alternative accounts of discriminative control, goal-directed responding does not spontaneously come under the control of the stimulus context in which the response-outcome contingency is experienced. Thrailkill and Bouton (2015) found that after limited instrumental training the magnitude of the devaluation effect shown by their rats was unaffected by a shift from the training context to another familiar context between the end of instrumental training and testing. It is unlikely that their rats did not discriminate between the contexts because with more extended training, when responding had become autonomous of outcome value (i.e., habitual), this context shift reduced overall responding. This pattern of results accords with our model in that after limited training the goal-directed strength *g* controls responding and, consequently, this control transfers spontaneously across contexts. By contrast, when responding has become under habitual control due to the monotonic increase in *h* with training, a context shift automatically produces a response decrement because such control reflects the development of context (stimulus)-response strength accrued during training.

As well as their paradoxical effect of extended training on outcome devaluation (see above), Garr et al. (2019) also reported an empirical evaluation of rate correlation theory by contrasting RI and fixed interval (FI) schedules. As well as replicating the established finding (Derusso *et al.*, 2010) that behavioural autonomy is more readily established by RI than by FI training, they also assessed the rate correlation within each training session. At variance with our account of goal-directed action, they report that, if anything, the FI rate correlation was lower than the RI rate correlation, even though FI was more sensitive to outcome devaluation. However, once it is acknowledged that goal-directed strength can come under discriminative control when the stimuli signal different rate correlations, the analysis of FI becomes much more complex. In the case of an FI schedule, the outcome itself functions as a discriminative stimulus signaling a period without any outcome. This discriminative function of the outcome is evident in the scalloped pattern of performance across the FI interval with low rates of responding immediately after the outcome and higher rates later in the FI. That the outcome served such a discriminative function in the Garr et al (2019) study is evident from the scalloped pattern of responding they report. As the extinction conditions in force during an outcome devaluation test are most similar to the stimulus conditions in force during the terminal period of the FI, our model predicts that the critical rate correlation that should be computed is that during the terminal period of the FI, which may well be higher than for a matched RI schedule. It is because of the complexities introduced by schedules in which the outcome has a discriminative role that we have restricted our empirical analysis to random or variable schedules in which this role is minimized^7^.

### Avoidance

We have developed rate correlation theory within a dual-system framework by reference to positive reinforcement of free-operant behavior using appetitive or attractive outcomes. However, Baum (1973) also analyzed free-operant avoidance in terms of his correlational Law of Effect. Under a typical free-operant avoidance contingency, a response causes the omission or postponement of a future scheduled outcome with the consequence that our recycling memory model yields a negative goal-directed strength (*g* < 0), at response rates that do not avoid all the schedule outcomes in a memory cycle. On the assumption that experience with the aversive outcome through incentive learning produce a negative incentive value, (*I* < 0), the product of the negative goal-directed strength and incentive value, *Ig*, will be positive and thereby contribute to the probability of a response being performed, *p*. Under our model, therefore, responding should increase under an avoidance procedure.

One of the most interesting aspects of avoidance behavior is that it requires an explanation of its persistence in the absence of an explicit reinforcing event (for a recent review, see Gillan *et al.*, 2016). In our model, once the response rate is sufficient to avoid all schedule outcomes within a memory cycle, the goal-direct strength will remain frozen at the established *g* value and thereby produce sustained avoidance in the absence of the aversive outcomes.

The most radical aspect of this account is its assumption that avoidance responding can be goal-directed. Although there are precedents for a goal-directed account of avoidance (e.g. Seligman *et al.*, 1973), contemporary RL theory follows traditional two-process theory in assuming that avoidance responding is purely habitual or model-free (see Maia, 2009). Although human discrete-trial procedures have demonstrated a reduction in avoidance following revaluation of the aversive outcome (Gillan *et al.*, 2011), more critical for a rate correlation account of goal-directed avoidance is a demonstration by Fernando et al. (2014) of an outcome revaluation effect using a free-operant schedule. They trained rats to lever-press to avoid foot-shocks that were programmed to be delivered at fixed intervals. Their revaluation procedure consisted of non-contingent presentations of the shock under morphine, so that pain would be reduced and the aversive status of the shock devalued. During an extinction test, their rats decreased responding compared to a non-revalued control group, demonstrating that their rats were performing the avoidance action to reduce the rate of an unpleasant outcome.

In accord with our dual-system model, Fernando and colleagues (2014) also investigated the role habit learning in free-operant avoidance. As mentioned above, an enduring problem for reinforcement theory is the absence of any event following an avoidance response that could act as a reinforcer. However, Konorski and Miller (1928) discovered that performance of an avoidance response itself, or more strictly speaking the feedback stimuli generated by responding, functioned as a conditioned aversive inhibitor and, subsequently, Weisman and Litner (1969) reported that an explicit aversive inhibitor acted as a conditioned reinforcer of free-operant avoidance responding by rats. Taken together, these results suggest that habitual responding may be reinforced by the feedback stimuli generated by responding itself. In accord with this analysis, Fernando et al. (2014) found that avoidance responding by their rats was enhanced by the presence of an explicit feedback stimulus and, moreover, this enhancement appeared to be habitual. Although exposure to the feedback stimulus under morphine enhanced its reinforcing property, the enhancement was not evident in an outcome revaluation test. This finding led Fernando and colleagues to conclude that the responding generated by the presence of the explicit feedback stimulus was habitual.

In summary, free-operant avoidance, like its appetitive counterpart, is under joint control by goal-directed and habitual systems with the former reflecting rate correlation learning between the response and aversive outcome, and the latter reinforced by the aversive inhibitory property of response-generated feedback stimuli.

## Discussion

In this paper we have formalized a theory of goal-directed actions and habits that marshals two systems to generate responses rates under random free-operant contingencies. After discussing associative and RL theories based on outcome probability and reward prediction-error, we presented an alternative theory of goal-directed control according to which agents compute a correlation between rates of responding and rate of outcomes in a fixed working memory to represent the current causal relationship between an action and its outcomes.

The new rate correlation is then combined with the historic average correlation stored in long-term memory to generate a goal-directed strength. Furthermore, we also assume that habit strength is acquired in parallel with goal-directed control through a standard prediction-error associative learning process and that the strength generated by these two systems simply summate to determine the rate of responding. The strength of the goal-directed input determines the extent to which responding is sensitive to the current value of the outcome as assessed by the outcome revaluation test, whereas the responding that is autonomous of the current outcome reflects the current habit strength. By simulation, we demonstrated how the theory captures instrumental performance under ratio and interval schedules when outcome probabilities (or rates) are matched, how goal-directed control transitions to habitual behavior with extended training and why this transition occurs more rapidly under an interval than under a matched ratio schedule. The model also explains why responding under choice procedures tends to remain under goal-directed control in spite of the amount of training when different outcomes are employed for each of the choice responses. In all these cases, the actual instrumental contingency implemented by a schedule is filtered by the *experienced* rate correlation generated by the agent’s performance under the particular schedule which, together with additional training constraints, generates the goal-directed response strength.

An important feature of our model is that our agent’s perception of the actual contingency may well be different to the objective contingency established by the schedule; we do not propose an optimal biological agent whose goal is to estimate the actual rate correlation in the environment. Our agent simply acts on the environment and experiences a rate correlation in each memory cycle that is only affected by the stability and/or vigor of the response strength during training— two of the cardinal features of habitual behavior (Balleine, 2019a). In this sense, the habit system not only is capable of inhibiting the goal-directed system throughout extinction, but also interferes with the experienced rate correlation that the agent computes in each cycle during training.

There are a plethora of formal accounts of free-operant performance (e.g. Killeen, 1994; Niv *et al.*, 2007; Peele *et al.*, 1984; Tanno and Silberberg, 2012; Wearden and Clark, 1988), at least one of which (Killeen, 1994) also deploys a short-term memory to couple recent behavior with reinforcement. However, none of these theories distinguish between goal-directed and habitual responding, which is the prime focus of our model, and so we have restricted our discussion to theories that recognize and implement this distinction. Apart from model-based and model-free RL, which we have have already discussed, we know of only one other theory that addresses the distinction within the context of free-operant behavior. Recently, Miller et al. (2019) presented a dual-system or -controller account in which they assume that at each time step the goal-directed system selects a response rate and observes the resultant outcome rate as determined by the schedule feedback function, which in turn provides a basis for selecting a response rate by comparing it with the effort exerted by the agent under a particular reinforcement schedule. This responding concurrently engages the habit controller, which contributes to the habitization of performance. By simulation of their model, they present acquisition profiles generated by the goal-directed and habitual systems under a VI and VR schedule that are similar to the ones generated by our model (See Figure 5) with the ratio schedule initially producing greater goal-directed control than interval training and with performance coming under habitual control with more extended training. However, their ratio-interval contrast is not based on matching the outcome probability across schedules, which, as we have seen, is a cardinal aspect for contrasting the effects of ratio and interval contingencies on responding. Moreover, they do not report whether their model can capture the impact of the primary variables of free-operant schedules, outcome probability, rate, and delay, nor whether their model anticipates the preservation of goal-directed control under choice or concurrent training with different outcomes.

A unique feature of the Miller et al.’s (2019) model is their account of habit learning which eschews the predominant view in the classic learning theory and RL literature that habits are reinforced by the contingent outcome. Instead, they resurrect Guthrie’s theory (1959) that simple stimulus-response contiguity is sufficient for habit learning. It is far from clear, however, that unlike Guthrie’s theory, their contiguity process can account for the extinction or reduction of an established habit. They illustrate the persistence of habitual responding by simulating an omission contingency by reference to a study reported by Dickinson, Squire, Varga and Smith (1998). Having extensively trained their model under positive contingency, Miller et al. reversed the contingency so that the outcome now only occurred in the absence of a response. When the response probability was close to one at the end of training, the imposition of the omission contingency had no impact on the probability of responding generated by their habit system. However, this simple contingency reversal does not simulate the actual procedure used by Dickinson et al. (1998). To detect the impact of the omission contingency, Dickinson et al. maintained reinforcement of the target response with the training outcome and superimposed an omission contingency with a different outcome. We know of no case in which habitual free-operant responding persists in the face of simple extinction (see Dickinson *et al.*, 1995), let alone an omission contingency. Simple stimulus-response contiguity is a nonstarter as a learning condition for free-operant habits.

Our model deploys a standard associative account of habit extinction based on the negative reward prediction-error resulting from the omission of an expected outcome. Importantly, this prediction error encodes the total prediction of both behavioral systems, which enables our model to provide a unique explanation of the survival of goal-directed control across extinction. As the goal-directed system can only compute a rate correlation when both responses and outcomes are represented in the memory, successful behavioral extinction requires inhibition of the historic goal-directed strength when the outcome is suspended. As a consequence, responding extinguishes because the sum of the strengths of the systems approaches zero, even though the goal-directed strength *g* remains positive with the value of the last rate correlation experienced during the training phase.

A further contrast between our dual-system model and that of Miller et al. concerns the interaction between the goal-directed and habitual systems. Whereas the outputs of the two systems simply summate to determine the probability of responding in our model, Miller et al. (2019) argue for a weighted average of the system outputs with the weighting varying the goal-directed and habit strenghs. This is the type of arbitration analysis taken within the context of human discrete-trial choice paradigms and has become a major focus for both empirical and theoretical research (Cushman, 2013; Daw *et al.*, 2011; Gillan *et al.*, 2015; Kool *et al.*, 2016; Kool *et al.*, 2017; Lee *et al.*, 2014). Such arbitration processes typically allocate control on the basis of the reliability of the estimations or the costs and benefits delivered by each system, but we know of no evidence that such arbitration processes play a role in free-operant responding.

The dual-system approach offered here differs sharply from RL models in how the systems interact during training. In contrast to deploying an additional arbitrator to determine which system controls responding according to the reliability of the estimations of each system, in our model the contribution of each system to performance is given by its relative strength across training, and total performance follows from the linear sum of their strengths. As just discussed, the mnemonic mechanisms of our model filter the actual instrumental contingency established by the schedule’s feedback function to yield the *experienced* rate correlation, which is modulated by the increasing habit strength during training. Consequently, behavioral control follows from the proportion of responding that is explained by the goal-directed system under different experimental conditions, not from an additional system that assigns control to either of the systems according to an additional computational process.

Another model that is relevant for our theory is one offered by Balleine and Dezfouli (2019; 2012), who argue that the functional unit in a behavioral stream may not be a single isolated action such as a lever press, but a habitual chunk generated by a chain of stimulus-response associations with each response in the chain functioning as a stimulus for the subsequent response. Within the context of free-operant responding by rats, the example they offer is the chain involving a lever press followed by an approach to the source of the outcome. In favor of this view of habits they cite an analysis of lever pressing for a food outcome by Halbout and colleagues (2019) who found that isolated lever presses were sensitive to outcome devaluation, and therefore goal-directed, whereas press-source approach chains were habitual. Although these results accord with the idea that free-operant response chains may be habitual, they do not speak to the issue of non-chunked lever pressing that is habitual, which is the main concern of our model. Therefore, we argue that a summation mechanism remains the simplest and most plausible account of non-chained actions established by free-operant contingencies.

Finally, we should acknowledge that a simple summation process has implications for the nature of the representations underlying instrumental learning. So far, we have focused on the learning process underlying the acquisition of instrumental knowledge while remaining agnostic about the nature of this knowledge (at least in the case of goal-directed action). Ever since Thorndike (1911) formulated his Law of Effect, psychologists have assumed that habit strength is implemented by excitatory (and inhibitory) stimulus-response associations, an assumption we have endorsed in our theory. Therefore, if we are to deploy simple summation between systems outputs, we have to assume that the goal-direct system also yields associative excitation (or inhibition). Appealing to such an associative output may not seem problematic, and indeed it has been widely assumed that goal-directed action is based on response-outcome associations. However, such isolated associations fail to capture the purposive nature of such actions, which require an appeal to representations with intentional properties, such as beliefs and desires (Dickinson, 1980). To address this issue, one possibility is embedding the associative representation within a cognitive architecture that implements a process of practical inference to derive an action intention (Dickinson, 2012). From this perspective, a future theoretical goal is to integrate such a processing architecture with the dual-system learning outlined in this paper.

## Acknowledgments

This work was part of OP’s Ph.D. work in the Department of Psychology and the Behavioural and Clinical Neuroscience Institute at the University of Cambridge, funded by CONICYT. OP thanks Mike Aitken and Amy Milton for supervision, and Nathan Holmes, John O’Doherty, Fabian Soto, Tor Tarantola, Gonzalo Urcelay and Fred Westbrook for insightful discussions and comments on this work. We also thank Yutaka Kosaki for providing us with valuable data to inform our simulations. OP would also like to thank the NHS for all the support provided to him and his family during his Ph.D. studies.

## Footnotes

1 Although the very high response rates sometimes observed under ratio training suggest the differential reinforcement of short IRTs, these rates reflect the acquisition of a new functional response, such as response bursting—our assumption in this paper will be that responding is fully random, meaning that it follows a Bernoulli process with a fixed probability of a response being performed in each second.

2 Although the exact analytic form of the feedback function for interval schedules is still a matter of debate (see Baum, 1992), it is well accepted that this function needs to flatten once response rates attain a sufficiently high level so that all outcomes programmed by the schedule are collected. A widely-accepted form of this function is 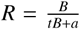, where *t* is the interval parameter and *a* is a parameter that depends on the conditions of the experiment, and which affect the animal’s pattern of responding independently of the outcome rate generated by the schedule.

3 In an open economy, the animal is also fed in the home cage with a different food to the one earned by the instrumental response during training, so that its weight remains constant throughout the experiment.

4 Previous versions of this algorithm deployed only one connection for increasing and decreasing the probability of responding. The original RL algorithm postulated by Bush & Mosteller (1951) had the form *h*_*t*+1_ = *h*_*t*_ + *α*^+^[1− (*h*_*t*_)] *α*^−^(*h*_*t*_) and assumed that *α*^+^ = 0 when a response was not reinforced. The term−*α*^−^(*h*_*t*_) can thus be regarded as reflecting an inhibitory potential present both in reinforced and non-reinforced responses.

5 The decrease in *p* under the value of *α*^−^ chosen for previous simulations made *p* decrease at a low rate and to remain at a positive and low value after 2000 memory cycles. Therefore, for illustrative purposes, we employed a higher value for *α*^−^ in the simulations shown in Figure 13 and kept the rest of the parameters identical to previous simulations (see Supplemental Material for the specific values used in the simulations presented in this paper).

6 This motivational effect of Pavlovian stimuli on instrumental responding is called *general* PIT, as it increases the probability of responding for all the available responses and is thought to be mediated by a general energizing effect of a stimulus that is associated with the motivational properties of the outcome. This is in contrast with *specific* PIT, where responding is enhanced only to the response that predicts the same outcome as in training and is thought to be mediated by the association between the stimulus and the sensory properties of the outcome (see Cartoni *et al.*, 2016 for a review).

7 In addition to manipulating the type of training schedule (RI vs FI), Garr et al. (2019) also varied the parameters of an RI schedule to generate a between-session rate correlation but found no effect of this variation on the outcome devaluation effect. However, our model computes the rate correlation locally, and Garr et al. report that the between-session variation in the RI parameter had no effect on the within-session rate correlation. Therefore, our model predicts that the between-session variation should also have been without effect on the magnitude of goal-directed control.

