## Supplemental Material for "A theory of actions and habits: The interaction of rate correlation and contiguity systems in free-operant behavior" (from another version of this preprint: version 1, 2019-10-17)

In this document we include multiple simulations that complement those shown in the paper. We also explain in detail how responses, reinforcement and experimental sessions were simulated and present the parameters used in the simulations for the dual-system model.

### General methods

All the following simulations were programmed in either Julia (julialang.org) or Python (www.python.org). Plots were produced by the seaborn package for Python. The relevant codes can be found online at github.com/omadav/dual_system_theory.

First, we explored the robustness of the correlation coefficient as a measure of the linear association between responses and outcomes in a free-operant setting across a considerable range of parametric variations. To this end, iresponse rate and sample length were used as independent variables. More formally, the simulations had the form $r = f(b, L, M)$, where $b$ is the response rate level, $L$ the length, or size of each sample and $M$ the length of the experiment. $r$ is, of course, the correlation coefficient that we were interested in investigating.

When the probability of observing a response in any given time is constant, response rate is an efficient measure of behaviour (Ferster & Skinner, 1957; Killeen, 1994). In this case, the distribution of IRTs will be best described by an exponential decay function. This is usually the pattern of responding observed under random schedules, which is the general purpose of our paper. Therefore, unless otherwise noted, in the following simulations responding was assumed to be completely random: although variation of responding is present at all times, we assume that the probability of a response to appear at each time-unit is fixed, so that at any given moment the occurrence of a response is completely unpredictable.

### Parametric investigation of a rate correlation theory

To study the the robustness of the correlation coefficient as a measure of the linear association between response and outcome rates, we modelled and simulated single experimental sessions. Each simulation represented one experimental subject under one particular schedule type and parameter (ratio requirement in the case of RR; interval requirement in the case of interval schedules). For each schedule and parameter, the process was replicated multiple times. Each session length was set to $M = mL$ sec, where $m$ is the number of time-samples and $L$ the length of each time-sample (in secs). For these simulations, the minimal unit of time was assumed to be $\frac{1}{3}$ of a sec. This condition effectively constrained subjects' responding to a maximum of 150 responses per min.

To register responses and outcomes in memory, a time-line composed of two binary vectors of the same length was created. Each vector represented the occurrence (represented by the number 1) and non-occurrence (represented by the number 0) on the responses and outcomes vectors. Such assumption made it possible to divide the time-line in $m$ samples and to consider the number of responses and outcomes in each. Finally, both responses and outcomes in each sample were aggregated and a standard correlation coefficient between these two-dimensional data points calculated.

A number of parameters were analysed for their effects upon the obtained correlations for the three different schedules of reinforcement of interest: RR, RI and RPI schedules. First, for each schedule and a given response or interval requirement, the effect of variations in time-sample length on the correlations was investigated, using a fixed level of responding. Second, for each schedule and a given response or interval requirement, the effect of the level of responding on the correlations obtained when the time-sample was kept constant was studied. The merits a rate-correlation theory will be evident if the correlation coefficient obtained is not seriously affected across a range of variations in these parameters. If that is so, then a correlational-based model of responding makes unambiguous predictions with regards to free-operant performance.

**Responding.** For each simulation, responding was simulated as follows: on each time-step, there was a fixed probability $q$ of a response to appear. The probability $q$ depended, naturally, on the number of responses per min ($B$) that we were interested in simulating:

$$P(resp = 1|B = b) = q = \frac{1}{3b} \quad (1)$$

where $b$ is the response rate simulated. For a baseline of, say, 60 responses per min, the probability of a response to appear on each time-step was fixed at $q = \frac{b}{3\times 60} = \frac{b}{180}$. The probability of responding on each second was therefore $p = 3q$.

**Reinforcement.**

**RR schedules.** A variable-ratio (VR) schedule provides reinforcement after a given number of responses, on average, have been recorded. The random-ratio schedule is an idealised version of this schedule. For each time unit, the RR schedule reinforces a response with a fixed probability. In a $RRk$ schedule, for example, the probability of each response being rewarded is $P(rnf|resp) = \frac{1}{k}$ every time a response is performed, where $rnf$ represents the event of a reinforcer being delivered. Both of these are binary variables taking values of 1 in their presence and 0 in their absence. To generate the RR schedule we note first that if the probability of a response being reinforced is $p$, then each time-unit can be regarded as a Bernoulli trial with $p$ as the probability of success, or the proability of a response being rewarded. Hence, each time a response occured, the reward value was drawn from a binomial distribution with parameters 1 and $p = \frac{1}{k}$.

**RI schedules.** On variable-interval (VI) schedules, a reinforcer is set up after an average time interval has elapsed. The first response observed after this interval is reinforced. RI schedules are an idealisation of VI schedules in that, for each moment, there is a fixed probability of a reinforcer being set up. Consequently, the generation of intervals in this schedule followed the same logic as that of responses in RR schedules; this time, however, the Bernoulli trial was in effect for each second that elapsed. On each second (3 time-units) the computer sampled either a 1 or 0 from a binomial distribution with size 1 and probability of success equal to the reciprocal of the preset interval. If this reward *flag* was equal to 1, then the next response was rewarded; once the reward was collected by the following response, the process started again. That is,

$$P(rnf|t) = \frac{1}{T} \tag{2}$$

where $t$ represents each second elapsed and $T$ the scheduled interval.

**RPI schedules.** For the RPI schedule, the probability of reward for the next response was calculated every time the simulated subject performed a response according to

$$P(rnf^*|resp) = \frac{x}{Tm} \tag{3}$$

where $rnf^*$ corresponds to the event of reward for the next response, $m$ is the memory size for the last $m$ IRTs (or $m + 1$ responses) and $x$ the time it took the subjects to perform the last m IRTs. Given Dawson & Dickinson's (1990) observation that $m$ does not have any systematic effects on performance, a value of $m = 20$ was used for the RPI simulations.

**Parametric manipulations.** For the RR schedule six ratio requirements were simulated: 5, 10, 20, 30, and 50. For RI and RPI simulations, the interval requirement was varied across 5, 15, 30 and 90 sec.

For all the simulations the sample length was varied across 10, 30, 60, 90 and 120 sec. Additionally, for every schedule and sample, 4 levels of responding (responses per min) were used. Level 1: 30 responses per min; Level 2: 60 responses per min; Level 3: 90 responses per min; Level 4: 150 responses per min. All the simulations were run with 300 samples and repeated 500 times. The values presented in the next section are averages of 500 data points for each of the conditions.

### Results

**RR schedules.** Figure 1 shows the results obtained for RR schedules. Inspection of the figure reveals that ratio schedules with lower ratio requirements generated consistently higher correlations than those with higher ratio requirements. The second observation that is relevant to the theory is that the time-sample length does not have any systematic effect on the correlations experienced in that there is no interaction between ratio requirement and sample length. This is an important theoretical point, as it allows a rate-correlation view of responding to make consistent predictions on performance under RR schedules: no matter what sample length any two subjects may be using, the subject trained with the lower ratio requirement should respond higher than that trained under the higher ratio requirement.

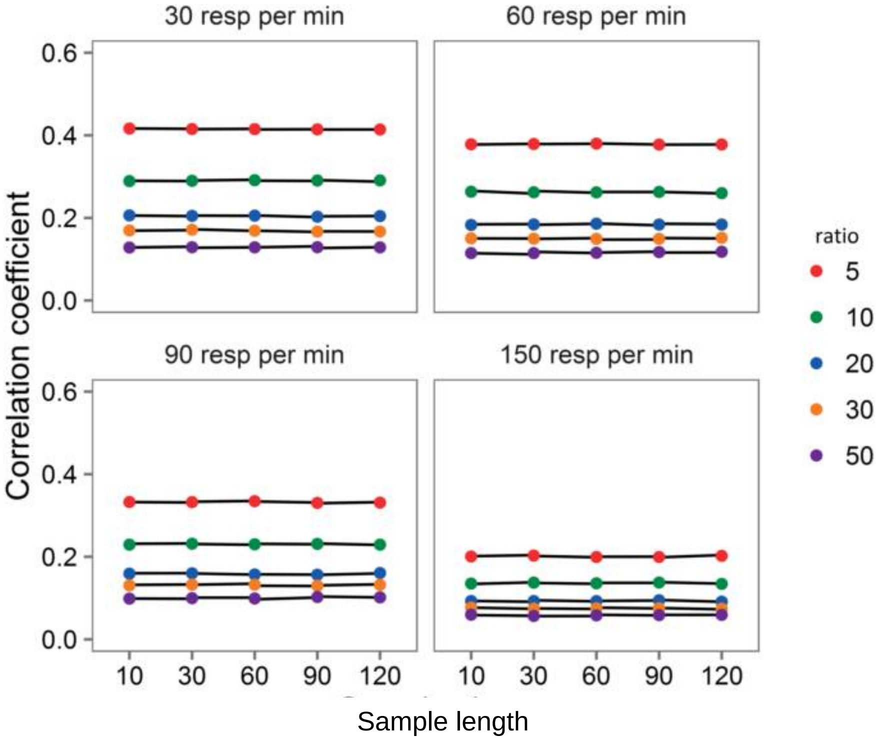


*Figure 1*. Correlation coefficient for RR schedules with different samples sizes (in sec). The legend signifies the ratio requirement.

**RI schedules.** Figure 2 shows the results obtained for RI schedules. Inspection of the figure indicates two main features of correlations under RI schedules that are similar to those observed in the case of RR schedules. First, with the exception of RI 5-sec and very low response rates, the effect of sample length on the correlation is very low for any given interval requirement. In addition, there is no interaction between sample length and interval requirement for any two different schedules. More importantly, however, is the observation that for the majority of response rates the correlations for RI schedules are close to zero. The observation is consistent with the slope of the feedback function for interval schedules, in that it flattens out after a certain level of responding; increasing response rates have no concomitant effects on outcome rate if the response rate is higher than the inverse of the interval requirement.

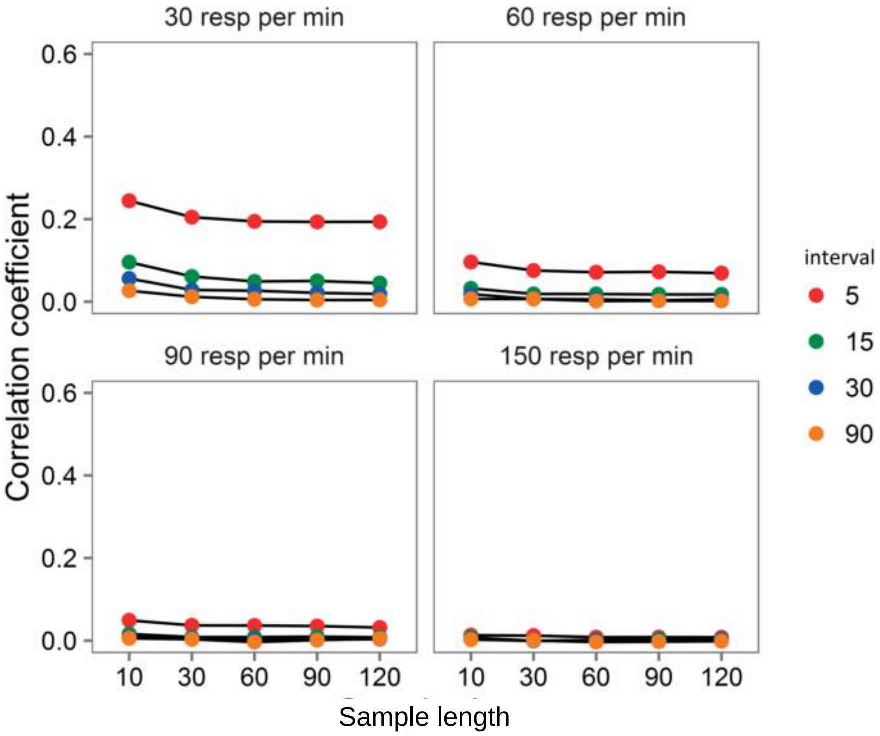


*Figure 2*. Correlation coefficient for RI schedules with different sample lengths (in sec). The legend signifies the ratio-equivalent value (in number of responses per reward).

In addition, similar to the RR case, the higher the interval requirement, the lower the experienced correlation. With increases in response rate, the difference in the obtained correlation converges to zero for all interval requirements.

At issue is the question of why a positive correlation for RI 5-sec and low response rates was obtained in these simulations. The explanation also follows Baum's feedback functions argument. Indeed, if the response rate is lower than the inverse of the interval requirement, then it is possible that increases in response rate will increase outcome rates. That is, to the extent that $b < \frac{1}{T}$, then increases in $b$ will be accompanied by increases in $r$. The feedback function for very low response rates then must have a positive slope. When the interval requirement is too low—or, in other words, the outcome rate is too high—it is even possible for the correlations to approach 1, making the RI effectively a FR1, or continuous reinforcement schedule.

An example may help illustrate the idea. Take the case of an RI 30-sec, so that the outcome rate is 2 outcomes per min. Suppose the subject is responding at a rate of 60 response per min, i.e., 1 response per sec. At this response rate, any increase in responding will not result in increases in outcome rates, since on average the subjects is effectively performing two responses per each reward. Suppose now that the rate of responding decreases to 30 responses per min, i.e., 0.5 responses per sec. At this new response rate, the subject is collecting all the outcomes with one response on average. However, if the subject slows responding to a level lower than this rate, then he/she will be losing otherwise obtainable outcomes. Increasing response rates now will increase the outcome rate; at the point where $b = \frac{1}{T}$, increases in response rate do not increase outcome rate. At lower rates, however, especially if the preset interval is short, increases in response rate will always be accompanied by an increase in the outcome rate and the correlation will approach 1. This particular feature of feedback functions for RI schedules has been extensively debated in the instrumental conditioning literature (see Baum (1992)).

**RPI schedules.** Figure 3 shows the results obtained for RPI schedules. The results are very similar to those for RI schedules. As expected, the experienced correlations are always lower for those RPI schedules with high interval requirements compared to those with low ratio requirements; the difference between these schedules tend to fade out as response rates increase. Additionally, for low response rates there is an effect of the sample length on the correlations: for the same interval requirement, increasing lengths lower the correlations.

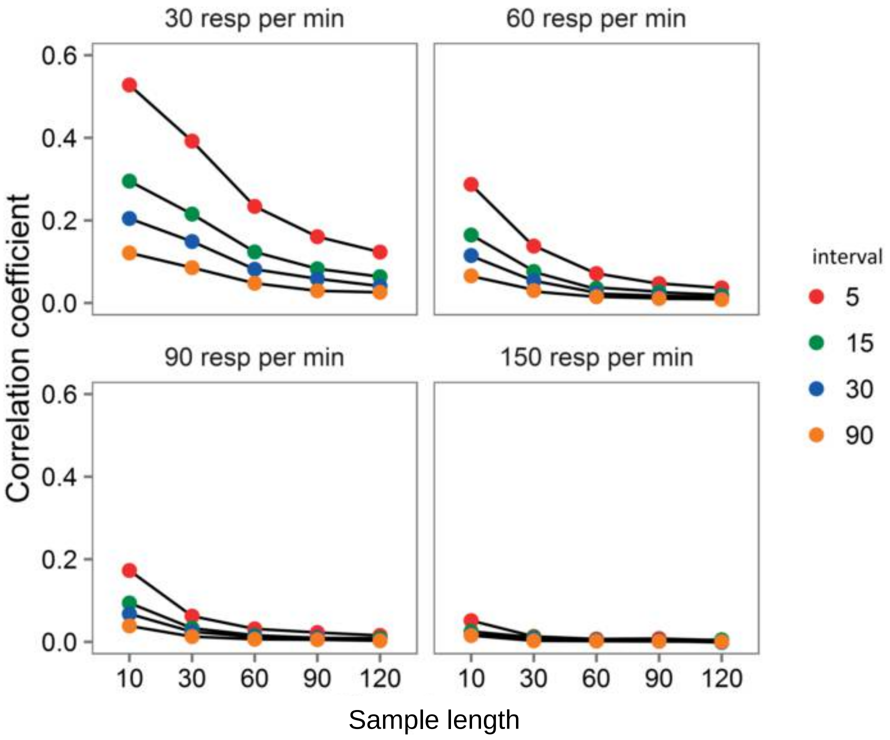


*Figure 3*. Correlation coefficient for RPI schedules with different sample lengths. The legend signifies the scheduled interval.

As in the RR and RI cases, the figure shows that there is no interaction between sample length and interval requirement on RPI schedules. Also evident from the RPI is the fact that RPI schedules with low interval requirements can generate positive correlations. In fact, contrasting this figure with that for RR schedules suggests that indeed these are comparable for some sample length values, especially the lowest ones.

**RI+ schedules.** The role of rate correlation as a determinant of instrumental performance has also found support in some studies with human participants using random interval plus-linear-feedback (RI+) schedules. In the RI+ schedule, the interval required for the *next* reward $I$ changes online according to

$$I = \frac{i}{n}b \tag{4}$$

where $i$ is the time since last reinforcement, $n$ is the number of responses performed during $i$ and $b$ is the equivalent ratio value. Thus, the probability of reward in each second will be changing according to the value of $I$, such that for a local response rate of $b_i = \frac{n}{i}$, the subject receives the reward on average after $b$ responses.

Figure 4 shows simulations of this type of schedule with the same parameters as those used for the RR simulations presented above. As can be seen in this figure, the RI+ schedule supports positive experienced correlations for a range of ratio-equivalent parameters. In addition, the RI+ schedule shares with RR schedules the property of lower correlations as response rates get considerably high. However, in contrast to RR schedules, the correlations increase when responding increases from a low level and start decreasing at sufficiently high rates.

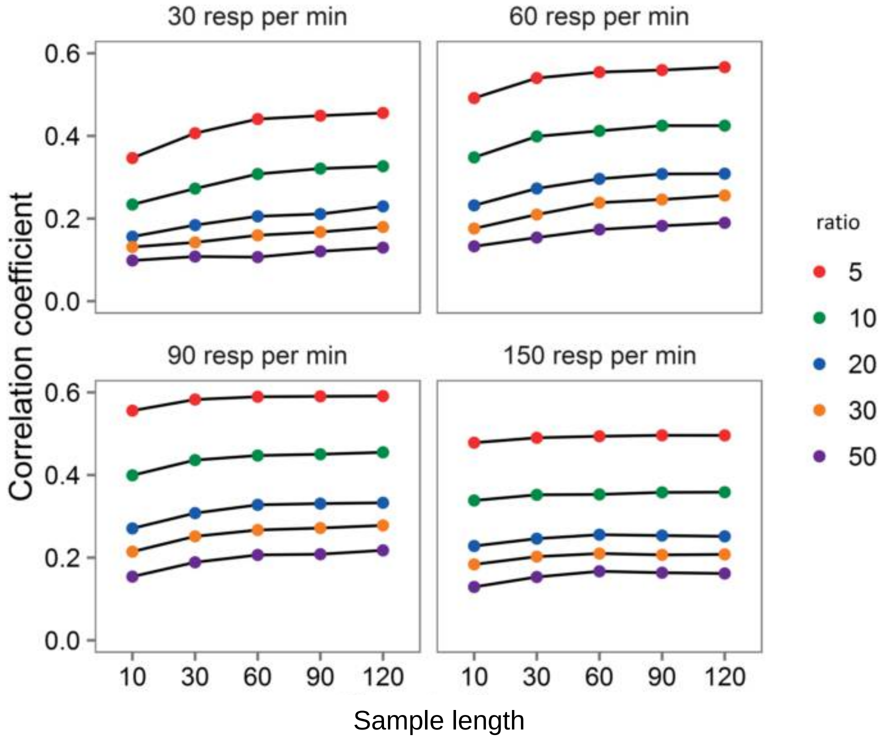


*Figure 4*. Correlation coefficient for RI+ schedules with different sample lengths (in sec). The legend signifies the scheduled interval (in sec).

### Parameters used in the simulations

As described in our paper, the dual-system model includes three main equations for different psychological processes, the habit system, the goal-directed system and the response activation function $s$ which transforms the total response strength $h + g$ into response probability per second. Each of these equations contains a number of parameters that need to be set. Given the wide range of experimental procedures and results across different species that nevertheless show the same directionality of effects, we show the generalizability of our model by choosing a suitable set of parameter values that captures the patterns of results across all of these experiments. We present the parameters for each of the sections in the table below.

| parameter | system | value |
|---|---|---|
| $\alpha^+$ | habit | $2 \times 10^{-2}$ |
| $\alpha^-$ | habit | $1 \times 10^{-5}$ |
| $\theta$ | goal-directed | .1 |
| C | response function | 0.6 |
| $\tau$ | response function | 10 |
